# Microbiome-induced Increases and Decreases in Bone Tissue Strength can be Initiated After Skeletal Maturity

**DOI:** 10.1101/2024.01.03.574074

**Authors:** C Liu, E.L. Cyphert, S.J. Stephen, B. Wang, A.L. Morales, J.C. Nixon, N.R. Natsoulas, M. Garcia, P. Blazquez Carmona, A.C. Vill, E.L. Donnelly, I.L. Brito, D. Vashishth, C.J. Hernandez

## Abstract

Recent studies in mice have indicated that the gut microbiome can regulate bone tissue strength. However, prior work involved modifications to the gut microbiome in growing animals and it is unclear if the same changes in the microbiome, applied later in life, would change matrix strength. Here we changed the composition of the gut microbiome before and/or after skeletal maturity (16 weeks of age) using oral antibiotics (ampicillin + neomycin). Male and female mice (n=143 total, n=12-17/group/sex) were allocated into five study groups:1) Unaltered, 2) Continuous (dosing 4-24 weeks of age), 3) Delayed (dosing only 16-24 weeks of age), 4) Initial (dosing 4-16 weeks of age, suspended at 16 weeks), and 5) Reconstituted (dosing from 4-16 weeks following by fecal microbiota transplant from Unaltered donors). Animals were euthanized at 24 weeks of age. In males, bone matrix strength in the femur was 25-35% less than expected from geometry in mice from the Continuous (p= 0.001), Delayed (p= 0.005), and Initial (p=0.040) groups as compared to Unaltered. Reconstitution of the gut microbiota, however, led to a bone matrix strength similar to Unaltered animals (p=0.929). In females, microbiome-induced changes in bone matrix strength followed the same trend as males but were not significantly different, demonstrating sex-related differences in the response of bone matrix to the gut microbiota. Minor differences in chemical composition of bone matrix were observed (Raman spectroscopy). Our findings indicate that microbiome-induced impairment of bone matrix in males can be initiated and/or reversed after skeletal maturity. The portion of the femoral cortical bone formed after skeletal maturity (16 weeks) is small; however, this suggests that microbiome-induced changes in bone matrix occur without osteoblast/osteoclast turnover using an, as of yet unidentified mechanism. These findings add to evidence that the mechanical properties of bone matrix can be altered in the adult skeleton.

## INTRODUCTION

Osteoporosis is a disease characterized by low bone mineral density and increased risk of fragility fracture. Although several effective pharmaceutical interventions are available to increase bone mineral density and reduce fracture risk, prolonged use of existing therapeutics either results in diminishing returns or increased risk of adverse side effects ^(1)^. Hence, new methods to address bone fragility are essential to advancing fracture prevention beyond current capabilities. The mechanical properties of bone tissue (the mineralized matrix) contribute to bone strength but are not directly addressed by existing therapeutics, although recent discoveries suggest the possibility of agents that enhance bone matrix^(2)^. Here we examine the gut microbiome’s effect on the strength of bone matrix.

The mammalian gut microbiome consists of bacteria, archaea, viruses, fungi and protozoa ^(3, 4)^. Changes in the constituents of the gut microbiome are associated with clinical conditions and ^(5)^can influence bone quantity and quality ^(6–10)^. We have previously shown that alterations to the composition of the gut microbiome can lead to impaired strength of cortical bone tissue ^(11–13)^. Specifically, when the composition of the gut microbiota is modified in mice through chronic dosing with specific types of antibiotics, the strength of cortical bone is reduced independent of geometry, indicating impairment of bone matrix.

Antibiotics are among the most powerful manipulations of the composition of the gut microbiome. Oral antibiotics rapidly change the composition of the gut microbiota by suppressing the growth of susceptible organisms^(14)^. In the days to months after an oral antibiotic regimen ends, the composition of the gut microbiota can return to that seen before treatment, although the time required for recovery can be lengthy ^(15)^. The composition of the gut microbiome can be rapidly reconstituted by applying a fecal microbiota transplant (FMT) from a donor ^(16)^.

There are two key limitations to prior studies examining the effects of the gut microbiome on bone matrix strength. First, our prior work manipulated the gut microbiota from weaning until euthanasia at skeletal maturity (16 weeks of life in mice)^(11, 12)^, a period when skeletal acquisition is most rapid and the majority of the bone volume within the cortical diaphysis is formed ^(17)^. It is therefore not clear if a change in the gut microbiota alters all the bone matrix or only regions of bone matrix formed after the microbiota is altered. If the gut microbiome only regulates bone tissue strength at the time of matrix synthesis, changes in the gut microbiota later in life (when bone formation rates are lower) are unlikely to lead to functionally important changes in whole bone strength, limiting the ability of the gut microbiota to address bone fragility in adults. Second, our prior work only examined male mice and it is unclear if changes in the gut microbiome have similar effects in females.

The goal of this line of investigation is to understand the effects of the gut microbiome on bone strength and fragility. Specifically in this study we address the following research questions: (a) How do changes in the composition of the gut microbiota applied after skeletal maturity influence bone tissue strength; and (b) how does the effect of the gut microbiome on bone tissue strength differ between males and females? To address these questions, we modified the gut microbiota in male and female mice after skeletal maturity by introducing oral antibiotics, removing oral antibiotics after skeletal maturity, or by reconstituting the gut microbiota using a fecal microbiota transplant (FMT). We hypothesize that the state of the gut microbiota alters bone tissue strength at the time of matrix formation, and therefore expect manipulation of the gut microbiota after skeletal maturity, when less bone volume is formed, to have only minor effects on bone matrix strength.

## 2.0 METHODS

### Animals

Animal procedures were approved by the local Institutional Animal Care and Use Committee. There can be considerable variation in the gut microbiota among animal shipments from the same vendor. To reduce variation in the gut microbiota among experimental groups, mice (C57BL/6J) were bred from a cohort of breeders obtained from a single vendor (Jackson Laboratory, Bar Harbor, ME, USA). Animals were housed in plastic cages with ¼-inch corn cob bedding (The Andersons’ Lab Bedding, Maumee, OH, USA), provided standard laboratory chow (Teklad LM-485, Envigo Diets, Madison, WI, USA), water *ad libitum*, a cardboard refuge environmental enrichment hut (Ketchum Manufacturing, Brockville, Canada), and raised in a 12:12 dark/light cycle. Mice were housed with their dam in a specific pathogen free facility until three weeks of age. At three weeks of age, the animals were randomly assigned cages of 4-5 animals separated by sex; cages were then moved to a conventional housing facility. To further reduce the effects of cage to cage variation in the gut microbiota within groups, soiled bedding was mixed between cages of animals from the same experimental group/sex each week until 12 weeks of age ^(18)^.

### Study Design

The study was designed to achieve a minimum of n=12 animals/group (power of 0.80 to detect an effect size of 0.88 with α = 0.05 using variance in bone tissue strength from prior work^(11)^). However, breeding was unexpectedly successful, and all available animals were used in the study. At weaning, pups from different breeding cages were placed in cages at random by sex. Later those cages were randomly assigned into 5 treatment groups per sex: Unaltered (n = 13 M, n = 14 F), Continuous (n = 17 M, n = 14 F), Initial (n = 18 M, n = 18 F), Reconstituted (n =12 M, n = 12 F), and Delayed (n = 14 M, n = 16 F). Experimental mice were bred from the same cohort of breeders in three separate breeding rounds (two months then six months after the first).

Animals in the Unaltered group received standard drinking water. Dosing groups received antibiotics via drinking water (1 g/L ampicillin and 0.5 g/L neomycin)^(11)^, an intervention we have previously shown results in reductions in bone matrix strength^(11)^. Ampicillin and neomycin exhibit low oral bioavailability, hence the antibiotics are poorly distributed systemically and the effect of dosing is primarily due to changes in the constituents of the gut microbiota ^(19, 20)^. The Continuous group was dosed from 4 weeks of age until euthanasia. The Delayed group received standard water until 16 weeks of age at which point dosing began and continued until euthanasia. The Initial group was dosed from 4-16 weeks of age, animals then received only standard drinking water for the remainder of the experiment with no intervention to ensure recovery of the microbiota. The Reconstituted group was dosed from 4-16 weeks of age, at which point a fecal microbiota transplant from sex- and age-matched untreated mice was applied to rapidly repopulate the gut microbiota to that of an Unaltered animal (see below for methods) (Figure 1A). These experimental groups include two groups in the gut microbiota was normal during rapid bone growth from 4-16 weeks of age (Unaltered and Delayed), three in which the gut microbiota was altered from 4-16 weeks of age (Continuous, Initial and Reconstituted), and three in which the microbiota was changed after 16 weeks of age (Delayed, Initial and Reconstituted). Fluorescent bone formation markers were injected at 12-, 16- and 24- weeks of age to identify regions of bone formed at different stages of the experiment (Figure 2B, C). Fecal pellets were collected at 16 weeks of age (immediately before any change in antibiotic dosing), and one day prior to euthanasia. Animals were euthanized at 24 weeks of age via cardiac puncture under anesthesia. Serum, cecal content, femurs, tibiae were collected immediately after euthanasia and stored at −80 °C. Perigonadal fat pads were collected and weighed. Serum samples were sent to Biomarkers Core at Duke Molecular Physiology Institute for biomarkers quantification.

**Figure 1.**
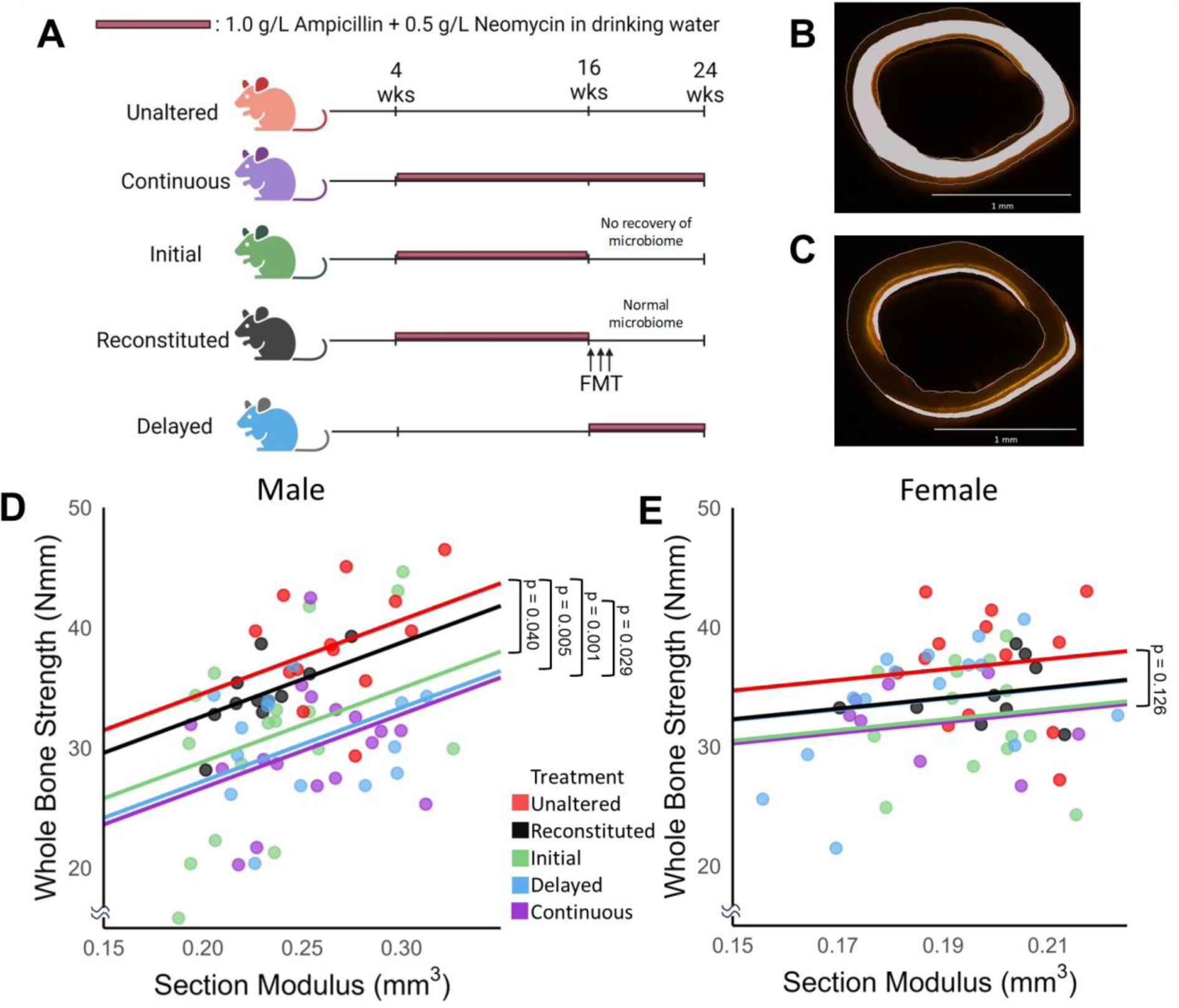
(A) Male and female mice were randomly divided into 5 experiment groups in which antibiotics were added to drinking water during different periods of life (highlighted). The mouse femur mid-diaphysis is shown with shaded regions indicating cortical bone formed during (B) 4-16 weeks of age and (C) 16-24 weeks of age as indicated by bone formation labels. (D, E) Graphical illustrations of the ANCOVA analysis comparing whole bone strength among groups after accounting for differences in section modulus are shown. The p-values indicate pairwise comparisons within the ANCOVA. In males, dosing before or after skeletal maturity impaired bone matrix strength, but Reconstitution of the gut microbiota restored bone matrix strength. Females showed similar trends but no significant differences among groups.

**Figure 2.**
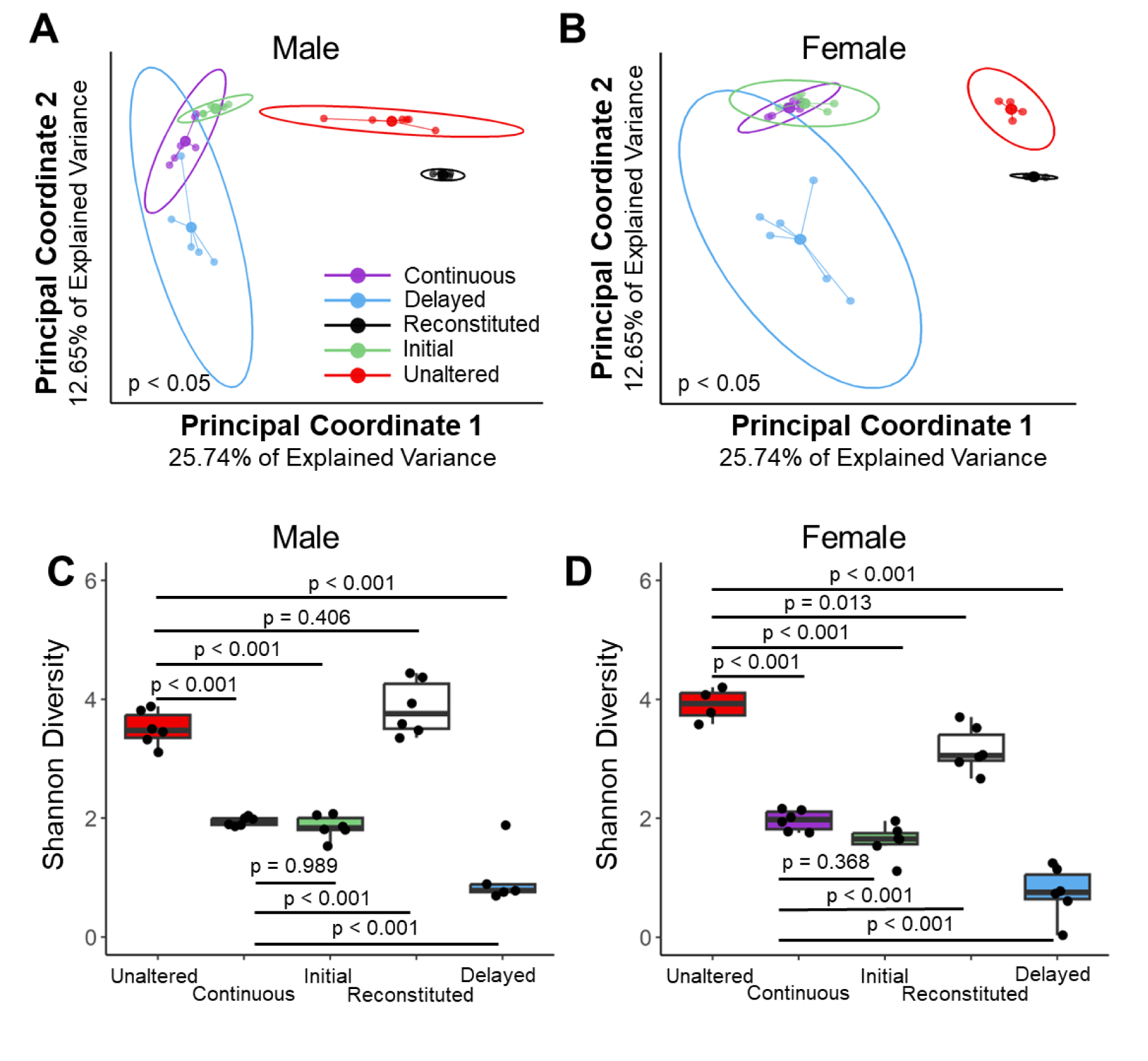
The composition of the gut microbiome for the five groups is shown for both sexes. The gut microbiome in animals receiving antibiotics at the time of euthanasia (Continuous, Delayed) was similar to that in the Initial group. The gut microbiome in the Reconstituted group was similar to that of the Unaltered group. The results are consistent with both beta diversity (male in A, female in B) and alpha diversity (male in C, female in D).

### Preparation of Fecal Microbiota Transplant

Fecal microbiota transplant was applied to animals in the Reconstituted groups as follows: Fresh stool samples (two pellets) collected from age- and sex-matched mice receiving standard drinking water were placed in sterile 1.5 mL microcentrifuge tubes. Fecal samples were immediately transferred to an anaerobic chamber (Coy Laboratory Products, Grass Lake, MI, US) and suspended in 1 mL anoxic PBS with 0.05% L-cysteine. Pellets were allowed to soften for 15 minutes and then vortexed for 3 minutes until the pellets were homogeneous. Suspended bacteria were separated from fibrous material by centrifuging for 5 minutes (150 rpm). Supernatant was aliquoted into a sterile bottle. Fecal slurries were pooled from multiple animals, one from males and one from females. The prepared bacterial solutions were placed in an anaerobic jar and transported to the animal facility. Mice were fasted for 3 hours prior to receiving the transplant (*ad libitum* access to water). Each mouse received 150 μL donor material from the corresponding age- and sex-matched pooled sample and placed in a fresh cage. The remaining slurry was frozen at −80°C for subsequent 16S rRNA sequencing. A fresh fecal transplant was applied to each mouse daily for three days. After fecal microbiota transplant, fecal samples were collected every two weeks to monitor recovery of the gut microbiota.

### Femoral Geometry of Mid-Diaphyseal Cortical Bone

The femurs were harvested, wrapped in PBS-soaked gauze and plastic wrap, and stored at −80°C prior to analysis. Images of the femoral diaphyseal cross section were obtained by microcomputed tomography using Bruker Skyscan’s Al + Cu filter (Skyscan 1276; Bruker, Billerica, MA; 100kV, 200µA) with a voxel size of 10 µm. The raw projection data was reconstructed using NRecon software (Bruker, Billerica, MA). Images were processed with custom script that uses a Gaussian filter (radius = 2) to remove noise and is then thresholded using the Otsu method. Femoral cross-sectional geometry was determined using 2.5% of the total bone length at the center, midway between the greater trochanter and lateral condyle (BoneJ, styloid-r11)^(21)^. The total area, cortical cross-sectional area, cortical thickness, neutral axis, moment of inertia about the medial-lateral axis were determined ^(22)^. The section modulus was calculated as the moment of inertia of the medial-lateral axis divided by the distance from the posterior surface to the neutral axis.

### Mechanical Characterization of Bone

The right femora were thawed to room temperature and remained hydrated during mechanical testing. Femur length was recorded using digital caliper as the distance between the greater trochanter to the lateral condyle. Femora were loaded to failure in three-point bending in the anterior-posterior direction at a rate of 0.01 mm/s using a span length of 7.5 mm between the supporting pins^(23)^ (858 Mini Bionix; MTS, Eden Prairie, MN, USA). Force and displacement were recorded using a 25lb load cell (MLP-25, Transducer Techniques, Temecula, CA, USA; accuracy confirmed through manual calibration each week). Whole bone strength was expressed as the maximum moment (half of the peak load multiplied by half of the span length)^(23)^. Differences in bone matrix strength between groups were detected as differences in whole bone strength after accounting for differences in section modulus using ANCOVA. This approach circumvents the assumptions made when calculating ultimate stress using beam theory assumptions ^(24)^. Additionally, tissue strength was calculated using the maximum moment divided by the section modulus and work to failure was calculated as the total area enclosed within the force-versus-deflection curve, spanning from zero deflection to the maximum deflection.

### Material Characterization using Raman Spectroscopy

After mechanical testing the proximal half of the femur was collected for Raman spectroscopy. The marrow was flushed out, the periosteum was removed and the sample was placed within. a Raman spectrometer equipped with a 785 nm red laser (Ramascope 2000, Renishaw, Mountain View, CA). Raman spectra were acquired at 50X magnification with 0.75 NA and 25% laser power (∼75 mW power), 15-second integration time and 5 accumulations. Spectral range was 380 cm^-1^ – 1800 cm^-1^, and grating was 1200 g/cm. The Raman microscope was calibrated against a silicon standard prior to imaging. Samples were scanned on the posterior side due to its relatively flat surface for better focusing; four point-Raman spectra were acquired sequentially from the proximal to distal regions for each sample. Given that the analysis was conducted on a partial femur specimen, the utilization of four point-spectra ensured optimal sample coverage without spectral overlap.

Cosmic ray removal was performed on all single spectra with the resident Renishaw analysis software. Background fluorescence removal and baseline correction were then performed on each individual spectrum using an adaptive fitting script in MATLAB (R2021b, MathWorks, Natick, MA). Corrected spectra were then averaged for each sample and processed in MATLAB to quantify peak intensity (I) and broadness (full width at half maximum, or FWHM). Peak broadness was used to calculate crystallinity (1/FWHM_960_), while peak intensities were used to calculate ν_1_PO4^3-^/Proline mineral-to-matrix ratio (∼I_960_/I_850_), ν_2_PO4^3-^/Amide III mineral-to-matrix ratio (∼I_440_/I_1240_), ν_1_PO4^3-^/Amide I mineral-to-matrix (∼I_960_/I_1660_), carbonate-to-phosphate ratio (∼I_1070_/I_960_), hydroxyproline-to-proline ratio (∼I_870_/I_850_), pentosidine (∼I_1495_/I_1450_), carboxymethyl-lysine (∼I_1150_/I_1450_), Amide I collagen disorganization (∼I_1660_/I_1640_), Amide I collagen conformational changes (∼I_1660_/I_1610_), and Amide I collagen maturity (∼I_1660_/I_1690_), as done previously^(25–32)^. Representative spectrum indicating the peaks and Amide I spectrum is shown in Supplemental Materials (Supplemental Figure 1).

To assess collagen quality and organization, the Amide I band was deconvolved first by identifying the position of four sub-peaks (∼1610 cm^-1^, ∼1640 cm^-1^, ∼1660 cm^-1^, ∼1690 cm^-1^) using a 2^nd^ derivative method as described previously^(27, 30–32)^. Only sub-peak positions identified within a 5 cm^-1^ window of the peak location were used. A fixed-position Gaussian curve fitting operation was then executed until a fit with R^2^ = 99% was achieved, and the resulting peak height of the deconvolved bands was used to calculate collagen disorganization, conformational change, and maturity. Both the 2^nd^ derivative sub-peak identification and peak fitting-deconvolution algorithm were performed in MATLAB using a script adapted from a freeware signal processing software (https://terpconnect.umd.edu/~toh/).

### Gut Microbiome Analysis

Microbial composition of fecal samples was analyzed using 16S rRNA amplicon sequencing. DNA extraction, purification, library preparation, and sequencing were performed by the University of California San Diego Microbiome Core utilizing previously published protocols ^(30)^. Samples were purified using the MagMAX Microbiome Ultra Nucleic Acid Isolation Kit (Thermo Fisher Scientific, USA) and automated on KingFisher Flex robots (Thermo Fisher Scientific, USA). Blank controls and mock communities (Zymo Research Corporation, USA) were included and carried through all downstream processing steps as quality control. 16S rRNA gene amplification was performed according to the Earth Microbiome Project protocol^(33)^. Illumina primers with unique forward primer barcodes^(34)^ were used to amplify the V4 region of the 16S rRNA gene (515fB-806r^(35)^). Amplification was performed as single reactions per sample^(36)^, then equal volumes of each amplicon were pooled, and the libraries were sequenced on the Illumina MiSeq sequencing platform with paired-end 150 bp cycles. QIIME2 (v. 2020.6) was used for quality control trimming and taxonomic classification using the SILVA database (SSU r138-1)^(37)^. Data was assigned at the genus level and normalized using a single rarefaction step with a feature count cut-off of 22287 (range of feature sizes in samples: 22401-130130; average and standard deviation feature size in samples: 69038 ± 22919)^(38)^. Standard rarefaction methods were used to determine feature count cut-off to maximize the number of features to percentage of samples retained^(38, 39)^.

The amplicon sequence variant (ASV) table generated by QIIME2 was used to calculate alpha (Shannon index) and beta diversity (Bray-Curtis dissimilarity) using the vegan package (v. 2.5-7) in R (v. 4.0.1)^(40)^. Principal coordinate analysis (PcoA) was performed on rarefied Bray-Curtis dissimilarity matrix and treatment group shown with 95% confidence level ellipses. Differentially abundant taxa at the genus level and above were compared among groups using a linear discriminant analysis of effect size (LefSe) (α = 0.05, LDA = 3)^(41)^. Additionally, a LefSe was performed to identify differentially abundant taxa between males and females. To reduce false discovery rate of microbial biomarkers, a combinatorial approach was used by performing Microbiome Multivariate Associations with Linear Models (MaAsLin2) to identify univariate associations between microbial abundance and treatment groups in conjunction with LefSe^(42)^. MaAsLin2 was performed including genera with > 10% prevalence (accounting for sex, treatment, and cage assignment) and post hoc q-values of the associations were calculated using the Benjamini-Hochberg method. A heat-map was generated depicting significant associations (q < 0.25) between genera and treatment groups relative to microbiota from the unaltered groups with their signed significance (sign of the correlation*-log10(q)). An association threshold of −5:5 was applied for visualization. *Prevotella* exceeded this threshold in all 4 treatment groups with an association of ∼-20. To evaluate engraftment and stability of the fecal microbiota transplant (taxa presence/absence) the unweighted Unifrac distance from baseline of Reconstituted mice relative to sex-matched fecal microbiota transplant donor slurry composition was determined using fecal pellets collected up to 8 weeks after the transfer ^(43–45)^.

### Statistical Analysis

Differences in the Alpha diversity (Shannon index) of the gut microbiome, fat pad weight, bone geometry, bone mechanical properties, and serum concentration measurements of different treatment groups were evaluated using a one-way analysis of variance (ANOVA). A post hoc Dunnett test was used to determine significance between different treatment groups relative to the Unaltered group. For parameters correlated with one another, ANCOVA, followed by a Tukey post hoc test, was employed to identify differences among groups after accounting for covariates. Similarly, characteristics of bone geometry were analyzed raw as well as after adjusting for animal body weight using a regression based approach^(46)^ Permutational multivariate analysis of variance (PERMANOVA) was used to determine differences between the Bray-Curtis beta diversity (microbiome composition) among treatment groups^(47)^. Unless explicitly specified, statistical tests were performed with a significance level of alpha = 0.05. Pearson’s product-moment correlation analysis was used to establish relationships between the Raman measurement and bone measurements^(48)^. All analyses were conducted using R Statistical Software (v 4.0.3; R Core Team 2020).

## Data Availability

Raw V4 16S rRNA DNA sequences are available at the NCBI’s Sequence Read Archive Database (BioProject ID: PRJNA1000601; http://www.ncbi.nlm.nih.gov/bioproject/1000601).

## 3.0 RESULTS

### Bone Geometry and Mechanical Properties

Males in the Continuous group had an impaired bone matrix strength that could not be explained by geometry showing a 35.2% reduction in whole bone strength compared to bones from Unaltered mice with similar section modulus (p = 0.001 ANCOVA), indicated graphically by the reduced intercept of the regression line in Fig. 1D. Similarly, whole bone strength was reduced in the Delayed (32.7% reduction, p=0.005) and Initial (25.5% reduction, p=0.040) groups compared to Unaltered mice with similar section modulus (Supplemental Table 1). The Reconstituted group exhibited a matrix strength similar to that of the Unaltered group (p = 0.929) and greater than that of Continuous group (p = 0.029). No significant differences in cortical bone cross-sectional area, moment of inertia, or section modulus of the mid-diaphyseal cortical bone were observed with or without adjustment for body weight (Supplemental Table 2).

In females, differences in bone matrix strength among groups followed a pattern similar to that seen in males, but no significant differences were detected (p =0.126 or greater, power = 51.7%, Fig. 1B). These findings suggest that if modification to the gut microbiota alters bone matrix strength in female mice, the effect is smaller than that seen in males and could not be detected with this sample size. No significant differences in cortical bone geometry (cross-sectional area, moment of inertia, section modulus) were observed among female groups either as measured or after adjustment for body weight (Supplemental Table 2.

### Changes in the Composition of the Gut Microbiota

In both males and females, the composition of the gut microbiome (Beta diversity) in Continuous, Delayed, and Initial groups was clustered together and differed from that in mice in the Unaltered or Reconstituted mice (Figure 2A). A PERMANOVA detected differences among treatment groups with males and females (p < 0.001). While there were differences in the composition of the microbiome between Unaltered and Reconstituted groups, the gut microbiota of the Reconstituted group clustered more closely to that of Unaltered mice (shifted to the right on the most influential principal coordinate). Both males and females demonstrated similar trends in Shannon diversity metrics among groups with Continuous, Initial, or Delayed groups having a significantly lower Shannon Diversity than the Unaltered or Reconstituted groups (Figure 2B; p < 0.001).

Fecal microbiota transplant changed the gut microbiota in the Reconstituted group; at 16 weeks (before removal of antibiotics) the composition was similar to that of the Initial group and Continuous group but at 24 weeks clustered close to that of the Unaltered group (see Figure 3A, Supplemental Figure 2, 3), demonstrating a modified microbiota before fecal microbiota transplant and recovery of the microbiota after transplant. In the Initial group, the composition of the gut microbiota resembled that of the Continuous group at both 16 and 24 weeks of age, indicating little recovery of the microbiota after 16 weeks, despite termination of antibiotic dosing. At 16 weeks of age the Delayed group resembled that of the Unaltered group, but after two months of dosing the microbiota resembled that of the Continuous group. Only small changes in the composition of the gut microbiota with time were observed in the Unaltered and Continuous groups.

**Figure 3:**
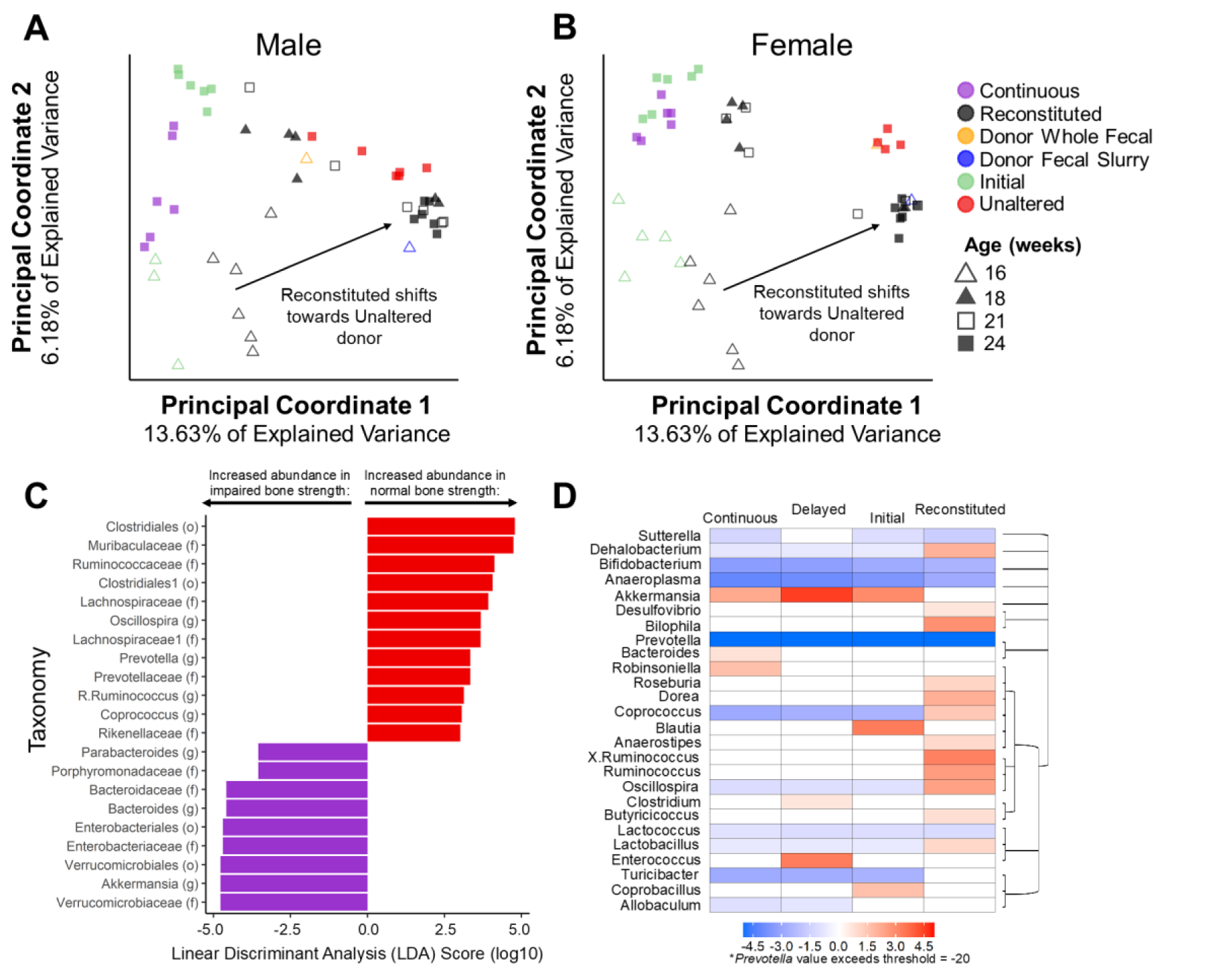
(A, B) Principal coordinate analysis of the microbiota of fecal samples collected at different time points between 16 to 24 weeks is shown. In the Reconstituted group, the composition of the microbiota shifted from that similar to the Continuous group at 16 weeks of age to one more similar to the Unaltered group at 24 weeks of age (shift to the right). Fecal pellets collected from donors (whole fecal) and processed for transplant (slurry) are similar to those of the Unaltered group. (C) The linear discriminant analysis of effect size comparing groups with impaired bone matrix strength (Continuous, Initial, Delayed) to groups with normal bone matrix strength (Unaltered and Reconstituted) is shown. Taxa are listed along with rank (f = family; g = genus, etc.). (D) The results of the Microbiome Multivariate Associations with Linear Models (MaAsLin) analysis are shown with taxonomies (relative to the Unaltered group).

A combinatorial approach of microbial biomarker analyses (LEfSe and MaAsLin2) was used to identify microbial features associated with a normal (Unaltered and Reconstituted groups) versus impaired bone matrix strength (Continuous, Delayed and Initial groups) in male mice at 24 weeks of age. The LEfSe analysis identified an increased abundance of *Bacteroides, Akkermansia*, and *Parabacteroides* and decreased abundance of *Oscillospira*, *Prevotella*, *Ruminococcus*, and *Coprococcus* in groups with impaired bone matrix strength (Continuous, Delayed, Initial) as compared to Unaltered and Reconstituted (Figure 3B). Animals in the Reconstituted group had increased abundance of *Oscillospira*, and *Ruminococcus* and decreased abundance of *Akkermansia*, *Bacteroides*, and *Prevotella* as compared to the Unaltered group. The MaAsLin analysis identified a strong positive association of *Akkermansia* (Continuous, Delayed, Initial groups) and *Enterococcus* (Delayed group) relative to the Unaltered group. Negative associations were identified between the *Lactobacillus* (Continuous, Delayed, Initial groups), *Lactococcus* (Continuous, Delayed, Initial, Reconstituted groups), *Coprococcus* (Continuous, Delayed, Initial groups), *Oscillospira* (Continuous, Delayed, Initial groups), *Turicibacter* (Continuous, Delayed, Initial groups), *Prevotella* (Continuous, Delayed, Initial, Reconstituted groups), and *Allobaculum* (Continuous, Delayed groups) relative to the Unaltered group (Figure 3B). Animals in the Reconstituted group had a strong positive association with *Desulfovibrio*, *Bilophila*, and genera from the *Bacillota* phylum including *Roseburia*, *Dorea*, *Coprococcus*, *Anaerostipes*, *Ruminococcus*, *Oscillospira*, *Butyricicoccus*, and *Lactobacillus* relative to the Unaltered group.

### Sexual Dysmorphism in the Composition of the Gut Microbiota

In males and females, the composition of the gut microbiome (Beta diversity) in the Continuous and Unaltered groups was comparable across sexes at 16 weeks of age (Figure 4A, p = 0.097 by sex, p < 0.001 by group). While there were significant differences in the composition of the microbiome between Unaltered and Continuous groups, the gut microbiota within the same treatment group was comparable between sexes. However, there were noticeable differences in the microbiome between Continuous males and Continuous females at 24 weeks of age (Figure 4B, p = 0.022 by sex, p < 0.001 by group). A LEfSe analysis was used to determine differentially abundant taxa between sexes in the Continuous group at 24 weeks of age (Figure 4C). The LEfSe analysis identified increased abundance of *Akkermansia* and *Robinsoniella* and a decreased abundance of *Enterococcus* and *Bacteroides* genera in the Continuous females relative to the Continuous males.

**Figure 4.**
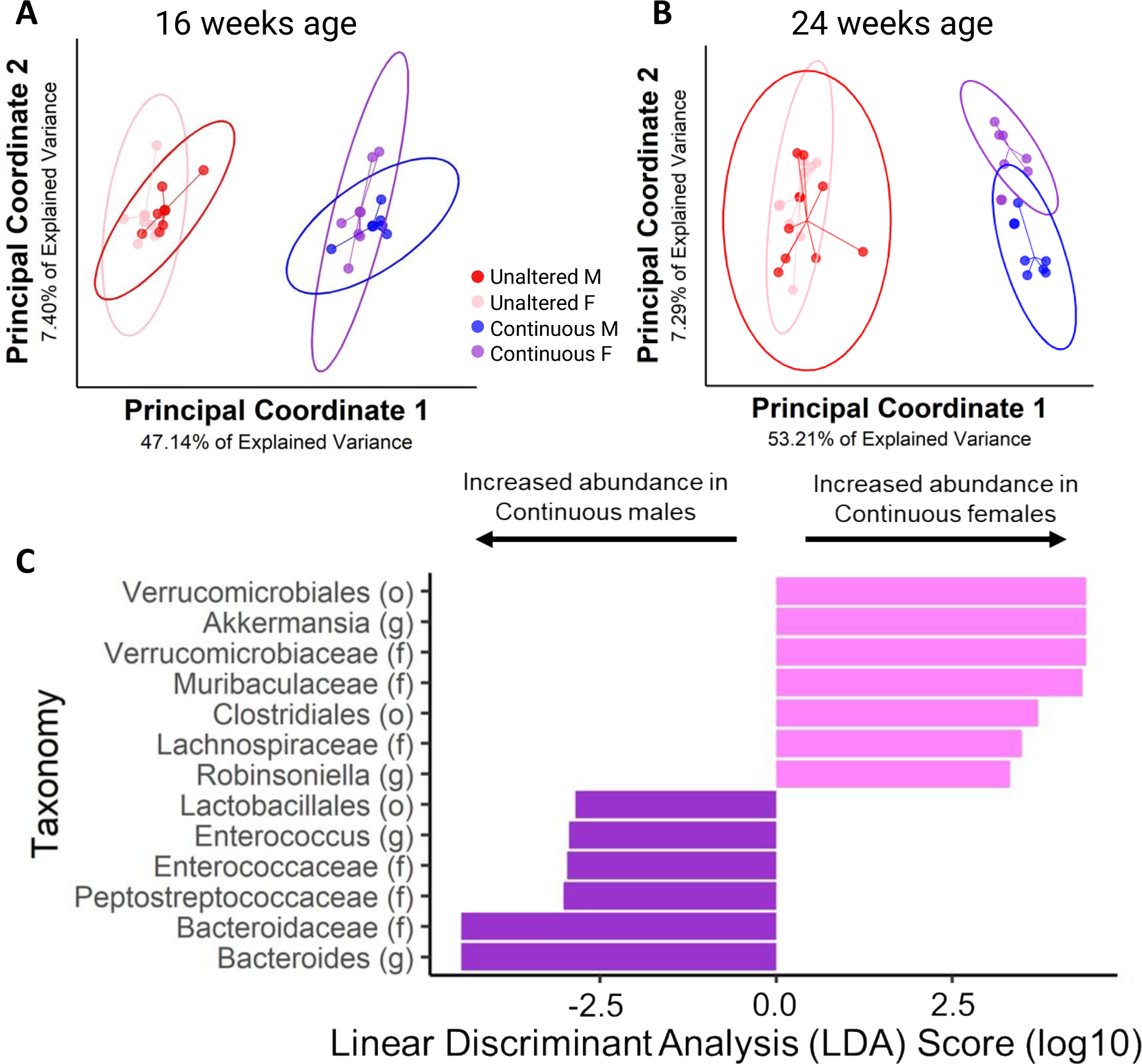
Differences in the gut microbiota between males and females are shown. Continuous antibiotic treatment exerted a differential sex-dependent effect on the composition of the microbiome during aging. (A) At 16 weeks of age no sex-dependent differences were apparent, but (B) at 24 weeks of age the antibiotics differentially influenced the microbiome of males and females. (C) Continuously treated females had an increased abundance of *Akkermansia* and *Robinsoniella* and decreased abundance of *Enterococcus* and *Bacteroides* relative to the males.

### Bone Tissue Composition

The composition of cortical bone tissue measured with Raman spectroscopy in the Reconstituted group was significantly different from that in the other groups (Figure 5). The mineral-to-matrix ratio, measured by the v1 phosphate peak (∼960 cm^-1^) to the amide I band (∼1600-1700 cm^-1^), was lower in the Reconstituted group when compared to the Unaltered group (p < 0.001) in males; the Initial group was lower compared to the Unaltered group (p = 0.036) in females. The averaged type-B carbonate substitution was decreased in the Reconstituted group compared to Unaltered (p < 0.001) in both males and females. Other Raman measurements that were not statistically different among treatment groups are presented in Supplemental materials (Supplemental Figure 4, 5).

**Figure 5.**
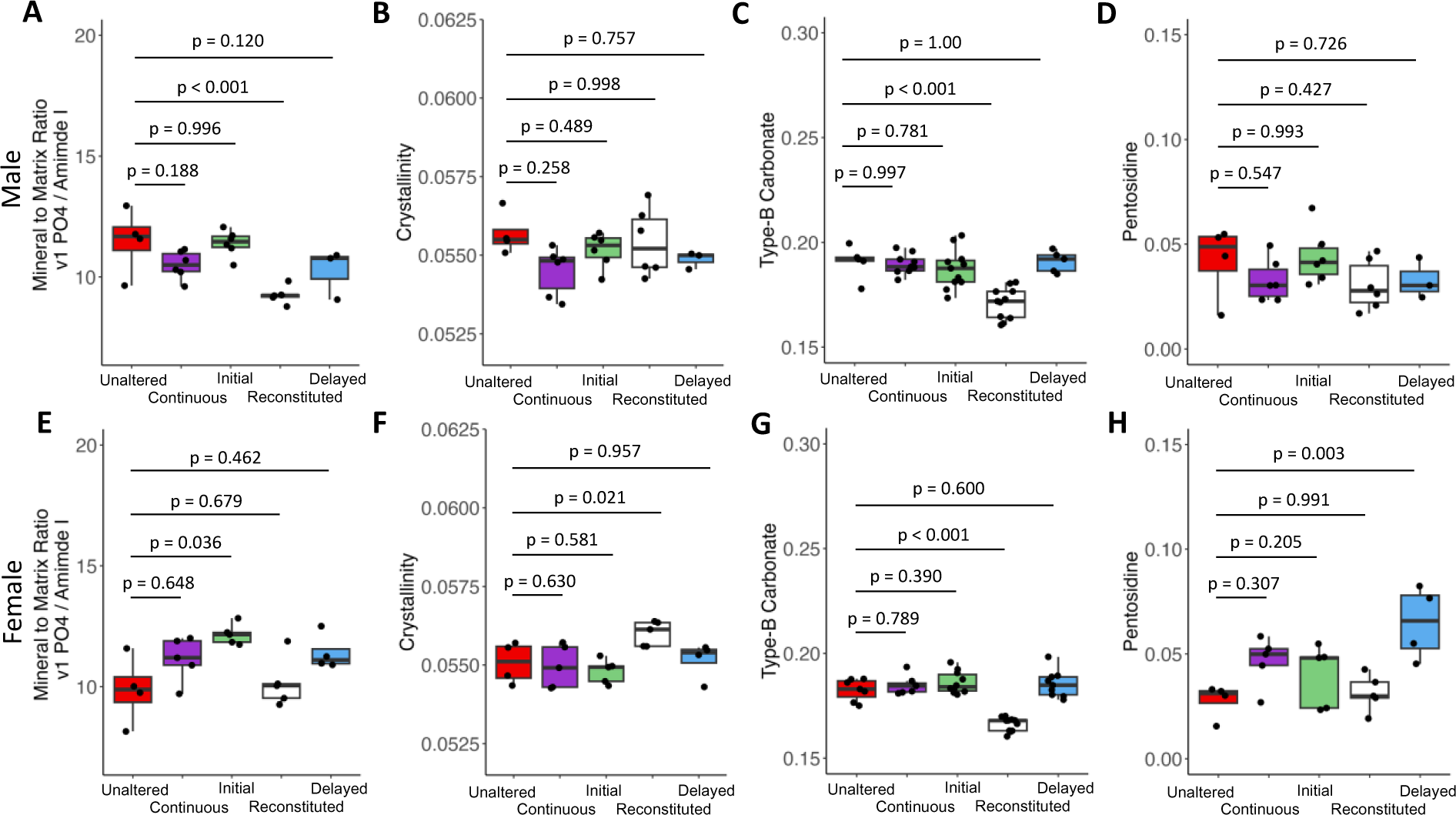
The chemical composition of bone matrix evaluated with Raman spectroscopy is shown. The mineral to matrix ratio, quantified by the area ratio of the v1 phosphate peak and amide I, demonstrated a reduction in Reconstituted males (A) but remained unchanged in females (E). Crystallinity did not differ across treatment groups in males (B) while it was increased in the Reconstituted females when compared to the Unaltered females (F). Type-B carbonate substitution was noticeably reduced in both Reconstituted males (C) and females (G) relative to the Unaltered groups. Assessment of Advanced Glycation End products (AGE) content, pentosidine measurements, indicated no changes across different treatment groups in the male (D) while increased pentosidine level was observed in the Delayed females (H).

In males, type-B carbonate substitution was negatively correlated with the tissue strength (r=- 0.362, 95% confidence level [-0.608 −0.052], p = 0.024, Supplemental Table 3), work to failure (r=-0.393, [-0.630 −0.088]), p = 0.013), and positively correlated with section modulus (r=0.447, [0.161 0.663], p = 0.003) and cross-sectional area (r=0.348, [0.046 0.593], p=0.026). The octacalcium phosphate-to hydroxyapatite ratio^(49, 50)^ was positively correlated with maximum moment (r=0.316, [-0.004 0.577], p=0.053) and cross-sectional area (r=0.341, [0.033 0.590], p=0.031).

In females, work to failure was negatively correlated to the mineral-to-matrix ratio (both the v1 phosphate peak to amide I band (r=-0.457, [-0.695 −0.129], p=0.009, Supplemental Table 4) and the v1 phosphate peak to proline peak (∼850 cm-1) (r=-0.372, [-0.631 −0.038], p=0.030), as well as type B carbonate (r=-0.547, [-0.750 −0.251], p=0.001). Amide I sub-peak ratios, such as the 1660/1690 ratio (enzymatic crosslink ratio^(51)^), were positively correlated with work to failure (r=0.479, [0.156 0.709], p=0.006); the 1660/1610 ratio (related to collagen conformational changes^(32)^) was negatively correlated to maximum moment (r=-0.432, [-0.675 −0.104], p=0.012), and strongly correlated to the tissue strength (r=-0.514, [-0.729 −0.208], p=0.002). The 1660/1610 ratio was positively correlated with cross-sectional area (r-0.492, [0.022 0.577], p=0.001) and section modulus (r=0.327, [0.022 0.577], p=0.037). Crystallinity was positively correlated with the section modulus (r=0.313, [0.010 0.564], p=0.043) and work to failure (r=0.479, [0.168 0.703], p=0.004).

### Serum Markers

The levels of serum procollagen I N-terminal propeptide (P1NP), a marker indicative of bone formation, exhibited distinct patterns across treatment groups. In males, the Reconstituted (p = 0.019) and Delayed (p = 0.021) groups showed lower levels compared to the Unaltered group. In females, the Continuous (p = 0.041) and Initial (p = 0.002) groups had higher levels of P1NP in comparison to the Unaltered group (Figure 6A, E). No significant differences in the serum tartrate-resistant acid phosphatase 5b were observed among groups in both male and female (Supplemental Figure 6B, E). Alterations to the gut microbiome did not lead to changes in serum Insulin-like Growth Factor 1 (IGF-1) (Supplemental Figure 6A, D). However, the proinflammatory cytokine Tumor Necrosis Factor Alpha (TNF-α) was significantly decreased in the Reconstituted groups when compared to the Unaltered groups in both males (p < 0.001) and females (p = 0.001) (Figure 6B, F). There were no significant changes in the adiponectin level among different treatment groups in males, but in females the Reconstituted group was reduced compared to the Unaltered groups (p = 0.011) (Figure 6C, G). In males, insulin levels were greater than the Continuous group as compared to Unaltered (p < 0.001) but no differences were detected among groups in females (Supplemental Figure 6C, F). No significant differences in leptin levels were observed among groups in males. In females, the Unaltered group had higher leptin levels compared to Continuous (p < 0.001), Initial (p = 0.009), Reconstituted (p = 0.017), and Delayed (p < 0.001) (Figure 6D, H).

**Figure 6.**
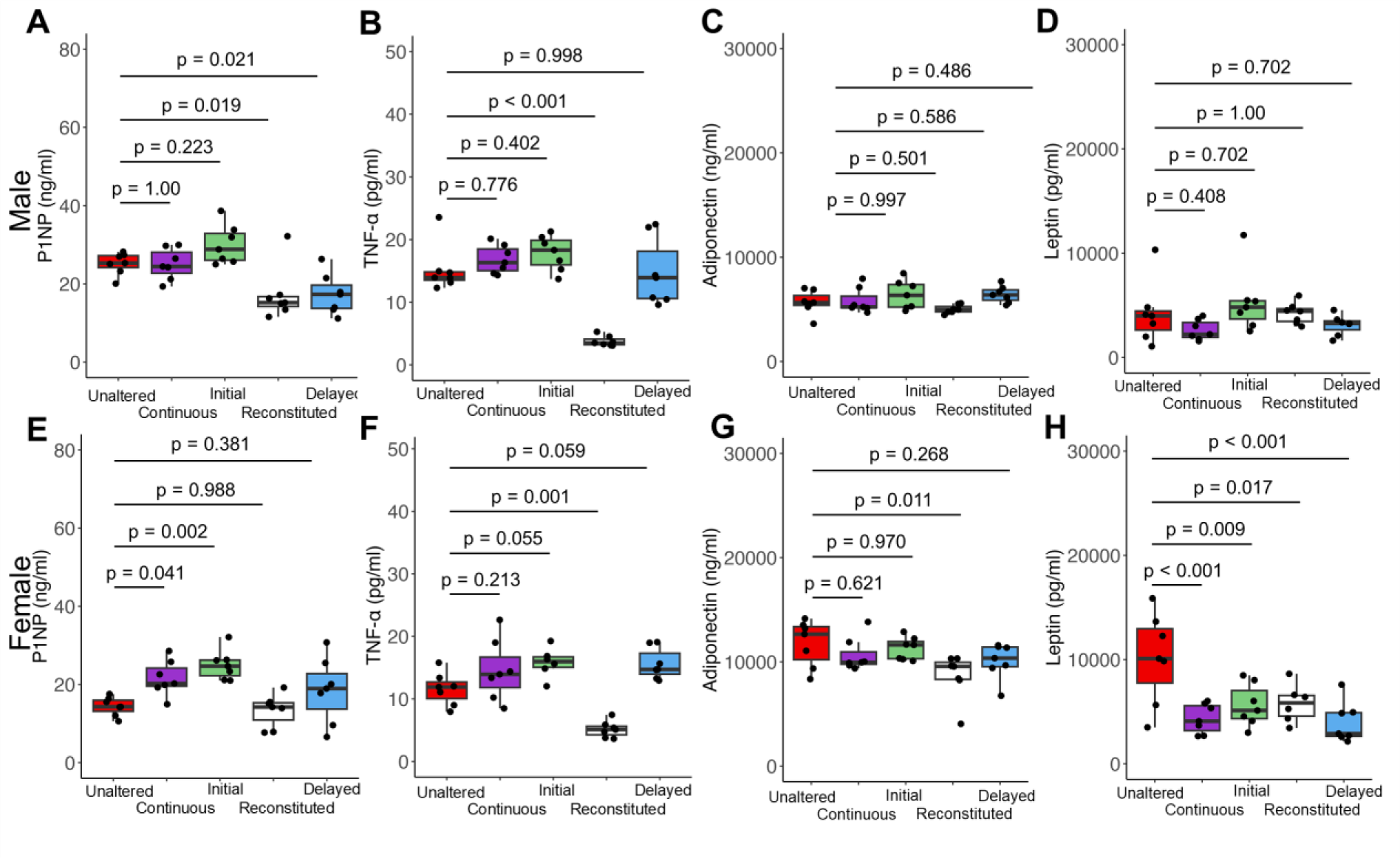
Serum markers are shown for (A, E) bone formation (P1NP); (B, F) TNF-α; (C, G) adiponectin; and (D, H) leptin for both sexes.

In males, the Interleukin 6 (IL-6), insulin, TNF-α, and adiponectin were correlated to several physiological measurements. IL-6 was positively correlated with the femur length (r=0.386, [0.037 0.651], p=0.032). Insulin was positively correlated with body weight (r=0.357, [0.027 0.617], p=0.035) and adiponectin was negatively correlated with body weight (r=-0.407, [-0.652 - 0.086], p=0.015). Serum TNF-α was negatively correlated with maximum bending moment (r=- 0.373, [-0.628 −0.046], p=0.027), body weight (r=-0.330, [-0.598 −0.003], p=0.053), and tissue strength (r=-0.401, [-0.648 0.078], p=0.017) (Supplemental Table 5).

In females, several serum markers were correlated with body weight and the work to failure (Supplemental table 6). In females, body weight was positively correlated to the level of adiponectin and leptin and negatively correlated to P1NP and TRAP5b. There were no detectable differences in the level of estrogen between treatment groups (Supplemental Figure 6).

## 4.0 DISCUSSION

In this study, we manipulated the constituents of the gut microbiota before or after skeletal maturity to determine the resulting changes in bone tissue. Our findings indicated that changes in the composition of the gut microbiome after skeletal maturity can decrease or improve bone matrix strength to the same degree as changes in the gut microbiota applied throughout growth and maturation. Since the amount of bone matrix formed after skeletal maturity is much smaller than that formed during growth^(17)^ (Figure 1B, C), our hypothesis that the gut microbiome influenced the composition of bone matrix only at the time of matrix synthesis is unlikely to be true. Furthermore, we showed that the effect of alterations to the gut microbiome on bone were greater in males than females.

Several lines of evidence support our finding that microbiome-induced changes in bone matrix strength in males were not limited to matrix formed after a change in the microbiota. First, tissue strength was altered when we applied changes in the gut microbiome after 16 weeks of age, when the bone formation rate within the cortex is greatly reduced ^(17, 52)^. If the gut microbiome exclusively influenced the properties of the matrix during bone formation, we would expect that the bone matrix strength in the Delayed group would resemble that of the Unaltered group, since only a small region of the femur cross-section was formed following the alteration of the gut microbiota (Figure 1C), however instead bone matrix strength in the Delayed group was more similar to that of the Continuous group. Similarly, bone matrix strength in the Reconstituted group was expected to be more similar to that in the Continuous group since the majority of bone matrix in the femur was formed while the gut microbiota was altered (Figure 1B), yet bone matrix strength in the Reconstituted group was more similar to that of the Unaltered group. In contrast, the Initial group displayed and gut microbiome and reduced bone strength more similar to the Continuous group despite removal of the oral antibiotics for the same period as the Reconstituted group.

One possible explanation for our findings was that mechanical failure of a whole bone under bending initiated at the regions of greatest tensile stress (in this case on the outmost posterior side), hence improvements in bone matrix strength only at the periosteal surface could have effects on measures of whole bone strength. However, bone tissue formed on the periosteum of the mouse femur after 16 weeks of age occurred primarily on the anterior side of the bone, not the posterior side of the bone^(17)^ (Figure 1B,C). Hence, changes in bone tissue in small regions of bone formed after 16 weeks of age could not explain the changes in bone matrix strength assessed here. Furthermore, serum markers of bone remodeling did not suggest that turnover of bone matrix after 16 weeks of age could explain the changes in bone matrix strength: the bone formation marker P1NP, was reduced in the groups in which the microbiome was changed after 16 weeks of age (Reconstituted and Delayed) and there were no differences in the bone resorption marker TRAP 5b among the treatment groups. Hence, histomorphometry and serum markers suggest that it is unlikely that there was substantial bone turnover after 16 weeks of age. Together, these findings demonstrated that an alteration in the gut microbiome changed bone matrix strength in as short as two months without substantial osteoclast/osteoblast turnover.

Our findings suggest that microbiome-induced changes in bone matrix strength could be explained by changes in bone matrix composition, although our findings are mixed. The strength of bone tissue is determined by the constituents of bone matrix including mineral content and crystallinity, collagen content and cross-linking, and the abundance of non-collagenous proteins^(53)^. Among the characteristics measured by Raman spectroscopy, the most convincing difference was a negative correlation between type B carbonate substitution and the mechanical performance of the bone (a finding also observed in other studies ^(54)^). We did observe a lower type B carbonate substitution in the Reconstituted group compared to all other groups, however type B carbonate substitution in the Unaltered group was not lower than that of the three groups with lower bone matrix strength (Continuous, Initial, Delayed), suggesting that type B carbonate substitution may not explain all of our findings. While we observed other significant differences in matrix composition using Raman spectroscopy, such differences were not consistent with observed differences in bone matrix strength, suggesting that the differences in bone matrix strength in our study were likely not due to changes in chemical composition detectable with Raman spectroscopy. A more comprehensive investigation is needed to explore other compositional changes in bone matrix that may be responsible for the observed effects on bone matrix strength.

Our findings highlight sex-related differences in response to antibiotic-induced manipulation of the gut microbiota. Microbiome-induced alterations in bone matrix strength were clear in males (25-35% effect), but were too small to be detected with this sample size in females (p = 0.126, power = 51.7%). A potential contributor to the differences between sexes was the response to dosing with ampicillin and neomycin. The composition of the gut microbiota in the Unaltered groups did not differ between males and females at either of the time points examined. However, Continuous groups (with the most drastic changes in the gut microbiota and bone) showed differences in the composition of the gut microbiota between males and females at 24 weeks of age. Hence, the microbiota in females appeared to respond differently to ampicillin and neomycin dosing, potentially explaining why the differences in bone matrix strength among female groups were more subtle.

The observed effects on bone matrix strength in males are likely due to changes in the microbiota, and unrelated to direct effects of antibiotics. First, the antibiotics used (ampicillin and neomycin) have low/zero oral bioavailability and therefore are not expected to be distributed to bone through the systemic circulation. Second, in the Initial group antibiotic dosing was suspended at 16 weeks of age, yet the microbiota and bone matrix strength at 24 weeks was similar to the Continuous group. If the antibiotics had a direct effect on bone, the matrix strength in the Initial group would have been more similar to that of the Unaltered and Reconstituted groups. This finding is consistent with our prior work which indicated that microbiome-induced changes in bone matrix strength could be caused by alterations to the gut microbiota caused by neomycin alone ^(12)^, yet when neomycin was dosed along with three other antibiotics causing removal of 99% of the microbiota bone matrix strength was not reduced as compared to Unaltered groups. These results suggest that microbiome-induced impairment of bone matrix strength depends on the specific microbial population which remains after dosing and not based on the exposure to neomycin alone.

Our analysis of the constituents of the fecal microbiota identifies several microbial taxa associated with changes in bone matrix strength. The five study groups showed two different phenotypes of bone matrix strength, normal bone matrix strength (Unaltered and Reconstituted) and impaired bone matrix strength (Initial, Delayed Continuous). Mice in the Reconstituted and Unaltered groups had increased abundance of genera from the *Bacillota (Firmicutes)* phylum relative to mice in the Initial, Delayed, and Continuous groups. This finding was corroborated across several metrics including relative abundance, LEfSe, and MaAsLin analyses. Both analyses consistently reveal significantly higher abundances and strong positive associations between the *Bacillota*: *Coprococcus*, *Lactobacillus*, *Oscillospira*, and *Ruminococcus*, in the Unaltered and Reconstituted groups as compared to the Initial, Delayed, and Continuous groups. Additionally, both analyses indicate increased abundance and strong positive associations between *Akkermansia* and *Bacteroides* in the Initial, Delayed, and Continuous groups compared to Unaltered and Reconstituted groups. Finally, the MaAsLin analysis identified a strong positive association of *Bilophila* and *Desulfovibrio* in the Reconstituted group relative to other treatment groups.

Although we did not measure microbial genes or metabolites, many of the genera with increased abundance in mice with normal bone strength are associated with two factors often associated with healthy gut microbiota: production of short chain fatty acids and modification of bile acids. *Oscillospira*, *Coprococcus*, and *Ruminococcus* produce the short chain fatty acid butyrate^(55–58)^. *Lactobacillus* produces lactic acid and L-Ornithine to maintain the gut mucosal barrier^(59)^ and elicits an immunomodulatory effect on bone^(60, 61)^. *Desulfovibrionaceae* and *Bilophila* produce hydrogen sulfide^(62, 63)^ which can mitigate bone loss by suppressing RANKL/OPG osteoclastogenesis ^(60, 64, 65)^. Furthermore, *Prevotella* (a producer of the short chain fatty acid propionate) showed increased abundance and were positively associated with the Unaltered and Reconstituted groups^(66–68)^. Mice in the Initial, Delayed, and Continuous groups had increased abundance of opportunistic microbes when the microbial diversity was depleted by dosing, including *Akkermansia* (all 3 groups), *Bacteroides* (Continuous group), and *Enterobacteriaceae* (Delayed group) ^(69–71)^. While these observations were notable, it remains unclear how these differences in microbial abundance may have led to an impaired matrix strength.

There were several limitations to our study. First, our hypothesis posited that the sexual dimorphism in bone phenotypes can be attributed to the dynamic changes in the gut microbiome that occurred between the ages of 16 and 24 weeks of age. However, the absence of mechanical bone data at 16 weeks precluded a direct assessment of whole bone strength at that specific age. While prior work in males shows dosing from 4-16 weeks of age causes reductions in bone matrix strength, without examination of bone matrix strength in both sexes at this age, we can only infer that the divergence in whole bone strength between sexes likely stemmed from differences in the composition of the gut microbiota. Second, three-point bending is not an optimal testing methodology for assessing bone tissue mechanical properties because bone cross-sectional geometry is irregular and the matrix is inhomogeneous ^(23)^ yet traditional calculation of matrix strength (maximum moment divided by section modulus) assumes the cross-sectional geometry is uniform and material composition is homogeneous. To avoid these limitations, we used ANCOVA to detect differences in whole bone strength that could not be explained by geometry. Hence, a more precise and micro-level mechanical testing approach is needed to accurately assess bone matrix strength.

Lastly, despite our comprehensive analysis encompassing bone composition, circulating bone turnover markers, hormones, and inflammation markers, the current study provides only limited insight into the factors that link the microbial taxa in the gut to bone or the specific changes in bone matrix caused by the microbiome. The challenge of identifying mechanistic links between the gut microbiome and organ phenotype remains the greatest challenge in the field of microbiome^(72)^. Understanding how the gut microbiota might alter the strength of bone matrix is further complicated by the fact that mechanisms regulating bone matrix strength are not as well studied compared to mechanisms that regulate bone volume/density^(73)^. Furthermore the mechanisms that are studied including nutritional deficiencies, genetic abnormalities or excessive tissue age, are not relevant to the current study in which diet, genotype and animal age were similar between groups^(13)^. To gain a deeper understanding, further investigations are required most likely using assessments of composition of the bone other than spectroscopy, measures of microbiota dependent metabolites in circulation and gene expression profiles of bone cells.

In summary, our findings demonstrate that the composition of the gut microbiome can influence the mechanical properties of bone matrix after skeletal maturity, suggesting that changes in the gut microbiota later in life (in adults) can alter bone matrix strength either reducing bone matrix strength to enhance bone fragility, or even improving bone matrix strength.

## ACKNOWLEDGEMENTS

This work was supported by NIH R01AG067997 (C.J.H.), NIH F32AG076244 (E.L.C.), and the Chan Zuckerberg Biohub. Dr. Teresa Porri for acquisition of micro-CT imaging data of the right femora. Janet Hueber for acquisition of the serum data. Imaging data was acquired through the Cornell Institute of Biotechnology’s Imaging Facility, with NIH S10OD025049 for the SkyScan 1276 mouse CT. We thank Mr. Raymond Dove for Nanoscale Characterization Core usage at Rensselaer Polytechnic Institute.

