## Supplemental Materials for "Microbiome-induced Increases and Decreases in Bone Tissue Strength can be Initiated After Skeletal Maturity"

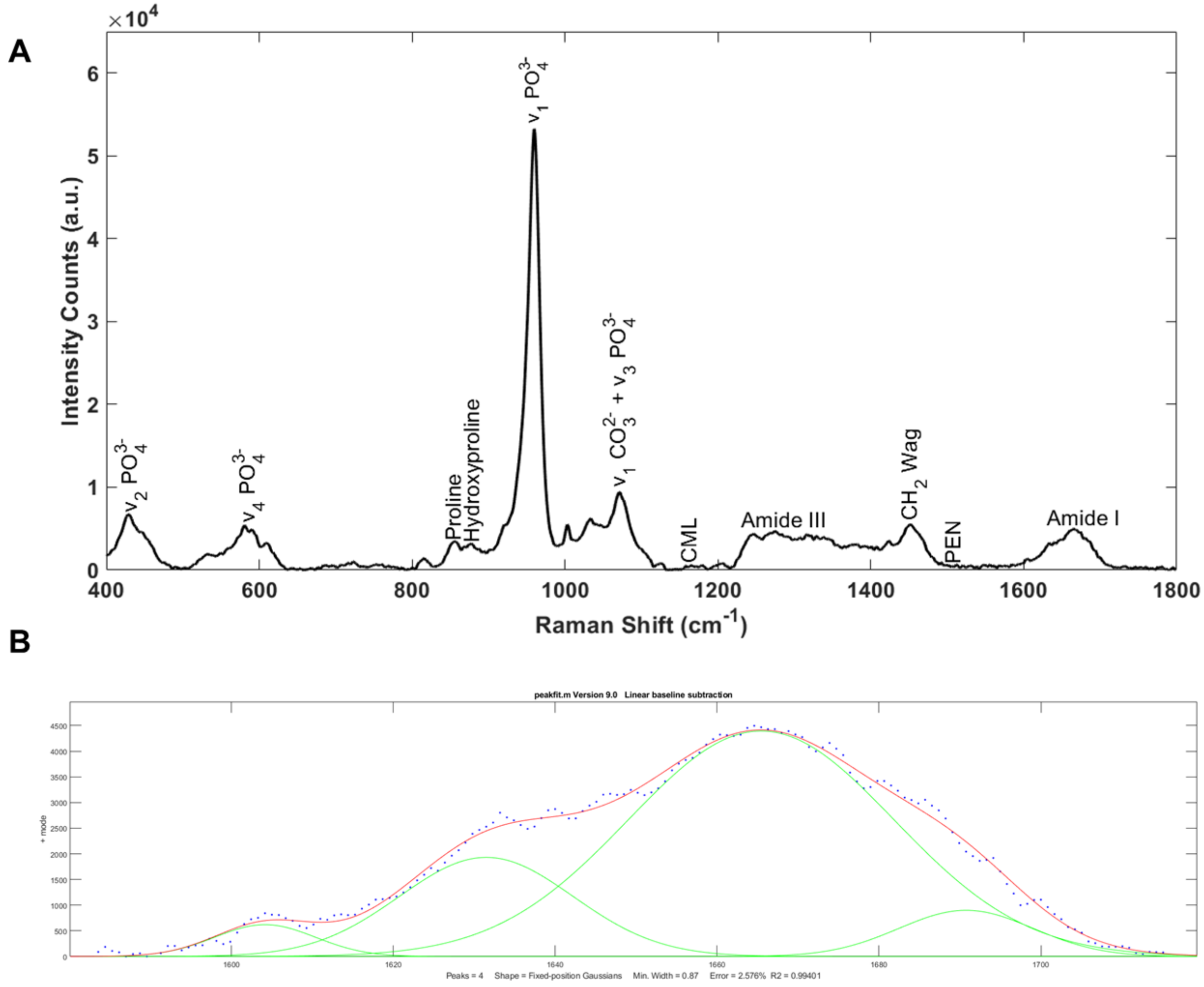


**Supplemental Figure 1.** (A) Representative spectrum indicating the peaks**.** (B) Deconvolution of the Amide I spectrum.

**Supplemental Table 1:** ANCOVA fit on generalized linear model.

| Parameter | Estimate | Standard Error | p-value |
|---|---|---|---|
| | ***Males*** | | |
| $a_0$, Intercept (Continuous) | 14.442 | 4.981 | 0.005 |
| $a_1$, Intercept (Delayed) | 0.548 | 1.880 | 0.771 |
| $a_2$, Intercept (Initial) | 2.162 | 1.816 | 0.238 |
| $a_3$, Intercept (Reconstituted) | 5.965 | 2.069 | 0.005 |
| $a_4$, Intercept (Unaltered) | 7.845 | 1.937 | < 0.001 |
| Slope | 61.172 | 18.914 | 0.002 |
| | ***Females*** | | |
| $a_0$, Intercept (Continuous) | 23.653 | 7.232 | 0.002 |
| $a_1$, Intercept (Delayed) | 1.965 | 1.878 | 0.300 |
| $a_2$, Intercept (Initial) | 0.243 | 1.962 | 0.902 |
| $a_3$, Intercept (Reconstituted) | 2.042 | 2.180 | 0.353 |
| $a_4$, Intercept (Unaltered) | 4.438 | 2.023 | 0.033 |
| Slope | 44.026 | 38.766 | 0.261 |

**Supplemental Table 2:** Biomechanics and femoral geometry metrics normalized by body weight. Comparison to sex-matched Unaltered group is shown (Dunnett test).

| Treatment Group | Femur length (mm) | Whole bone strength (N*mm) | Cross-sectional area ($mm^2$) | Moment of inertia ($mm^4$) | Section modulus ($mm^3$) | Tissue strength ($N/mm^2$) | Distance to neutral axis (mm) | Work to failure (N*mm) | Max force (N) |
|---|---|---|---|---|---|---|---|---|---|
| | | | | | ***Males*** | | | | |
| Unaltered | 16.199 ± 0.234 | 39.401 ± 4.604 | 1.018 ± 0.065 | 0.176 ± 0.020 | 0.262 ± 0.025 | 150.661 ± 16.028 | 0.669 ± 0.028 | 6.010 ± 3.907 | 21.014 ± 2.456 |
| Continuous | 16.169 ± 0.160 p=0.985 | 29.820 ± 4.813 p<0.001 | 1.003 ± 0.079 p=0.937 | 0.170 ± 0.023 p=0.846 | 0.254 ± 0.029 p=0.715 | 118.810 ± 23.359 p<0.001 | 0.669 ± 0.028 p=0.999 | 1.743 ± 1.497 p<0.001 | 15.904 ± 2.567 p<0.001 |
| Initial | 16.223 ± 0.278 p=0.993 | 32.735 ± 6.259 p=0.002 | 0.982 ± 0.082 p=0.413 | 0.166 ± 0.020 p=0.537 | 0.250 ± 0.024 p=0.453 | 131.410 ± 25.939 p=0.040 | 0.664 ± 0.025 p=0.962 | 2.747 ± 1.947 p<0.001 | 17.459 ± 3.338 p=0.002 |
| Reconstituted | 15.950 ± 0.153 p=0.019 | 34.479 ± 3.010 p=0.054 | 0.945 ± 0.060 p=0.043 | 0.143 ± 0.017 p<0.001 | 0.230 ± 0.019 p=0.004 | 150.214 ± 6.842 p=0.999 | 0.620 ± 0.030 p<0.001 | 5.661 ± 1.875 p=0.987 | 18.389 ± 1.605 p=0.054 |
| Delayed | 16.163 ± 0.193 p=0.977 | 30.462 ± 4.451 p<0.001 | 0.984 ± 0.050 p=0.521 | 0.178 ± 0.019 p=0.990 | 0.259 ± 0.016 p=0.993 | 117.745 ± 18.332 p<0.001 | 0.686 ± 0.037 p=0.393 | 2.372 ± 1.536 p<0.001 | 16.247 ± 2.374 p<0.001 |
| | | | | | ***Females*** | | | | |
| Unaltered | 15.726 ± 0.184 | 36.716 ± 4.854 | 0.867 ± 0.036 | 0.123 ± 0.008 | 0.195 ± 0.010 | 188.672 ± 27.993 | 0.632 ± 0.021 | 2.950 ± 1.314 | 19.582 ± 2.589 |
| Continuous | 15.699 ± 0.165 p=0.990 | 32.740 ± 2.907 p=0.098 | 0.849 ± 0.054 p=0.784 | 0.118 ± 0.013 p=0.501 | 0.191 ± 0.019 p=0.861 | 173.172 ± 24.524 p=0.395 | 0.617 ± 0.013 p=0.247 | 2.316 ± 0.924 p=0.478 | 17.461 ± 1.550 p=0.098 |
| Initial | 15.845 ± 0.148 p=0.239 | 32.936 ± 4.465 p=0.069 | 0.859 ± 0.042 p=0.979 | 0.124 ± 0.007 p=0.999 | 0.197 ± 0.011 p=0.996 | 168.162 ± 25.921 p=0.102 | 0.630 ± 0.015 p=0.999 | 2.600 ± 1.047 p=0.824 | 17.566 ± 2.381 p=0.069 |
| Reconstituted | 15.980 ± 0.195 p=0.005 | 34.652 ± 2.511 p=0.612 | 0.850 ± 0.024 p=0.806 | 0.124 ± 0.012 p=0.999 | 0.197 ± 0.012 p=0.997 | 176.361 ± 12.175 p=0.597 | 0.627 ± 0.029 p=0.953 | 5.088 ± 1.167 p<0.001 | 18.481 ± 1.339 p=0.612 |
| Delayed | 15.817 ± 0.175 p=0.452 | 34.473 ± 4.398 p=0.422 | 0.851 ± 0.054 p=0.749 | 0.121 ± 0.009 p=0.920 | 0.189 ± 0.012 p=0.570 | 182.430 ± 23.585 p=0.904 | 0.639 ± 0.017 p=0.708 | 2.584 ± 0.915 p=0.792 | 18.386 ± 2.345 p=0.422 |

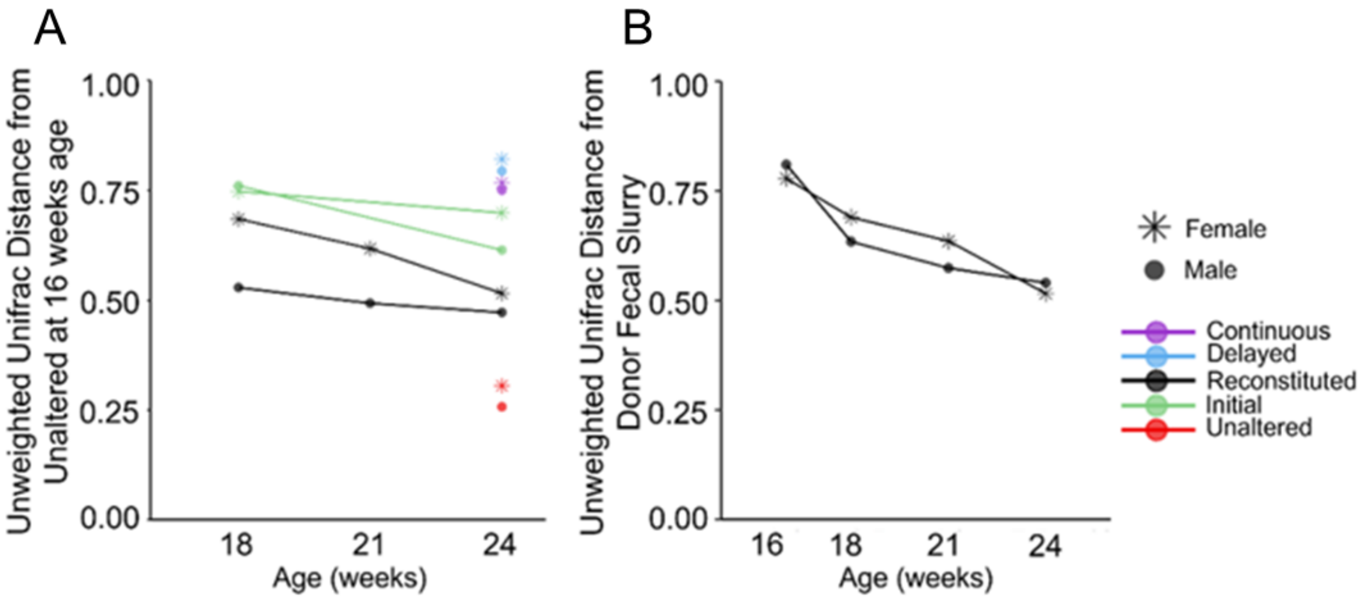


**Supplemental Figure 2.** (A) The fecal microbiota transplant successfully engrafts in the Reconstituted group and stabilizes towards the composition of the Unaltered group over an eight-week period, as compared to the Initial group which remains close similar to that of the Continuous group. (B) The microbital composition in the Reconstituted group shifts towards the composition of the Unaltered donor slurry.

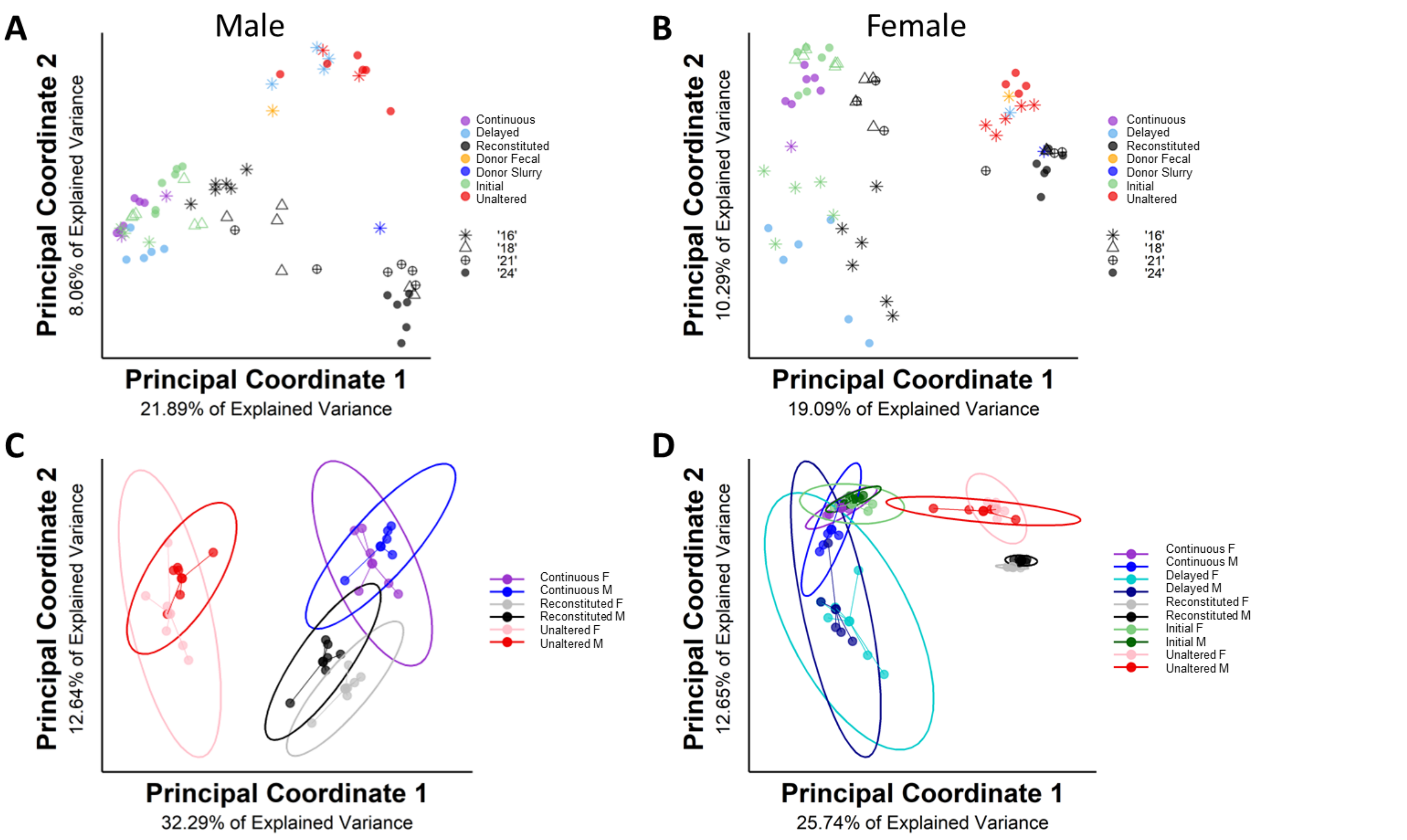


**Supplemental Figure 3.** (A,B) Principal coordinate analysis of fecal samples collected from 16 to 24 weeks reveals that the composition of the gut microbiota within a group remained relatively stable over time in the absence of a change in dosing in both males and females. (C) Combined male and female Bray-Curtis beta diversity analysis at 16 weeks age showed that the Reconstituted group have similar composition of the gut microbiome to the Continuous groups in both males and females. (D) Combined male and female Bray-Curtis beta diversity analysis at 24 weeks age demonstrated that for most treatment groups (other than Continuous and Reconstituted) males and females had a similar microbiota composition.

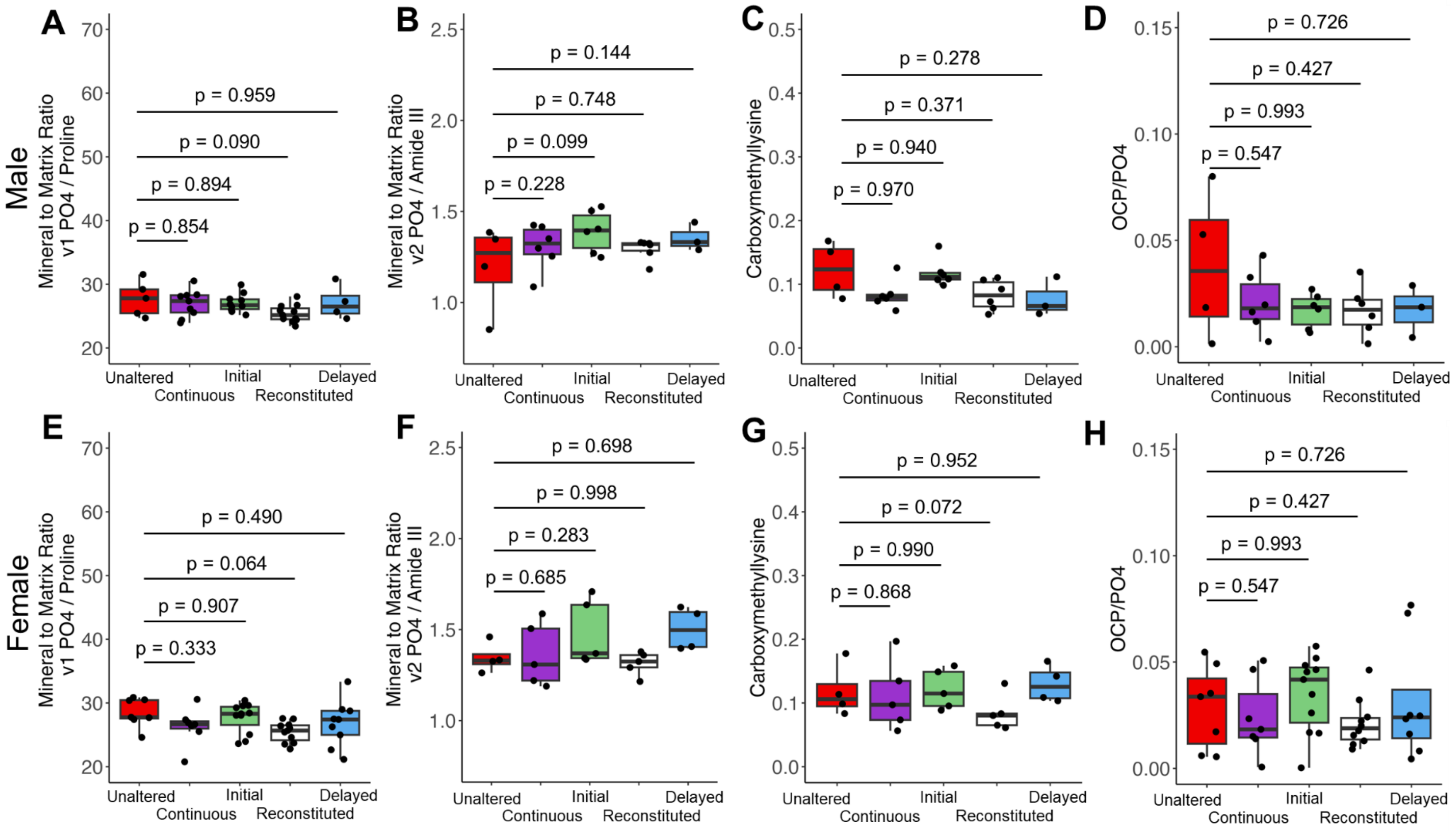


**Supplemental Figure 4.** Raman spectroscopy measurements of the mineral-to-matrix ratios, represented by the ratio of the v1 phosphate peak to proline (A, E) and the ratio of the v2 phosphate peak to the amide III peak (B, F), which did not differ among treatment groups. The advanced glycation end product (AGE) represented by Carboxymethyllysine (CML) (C, G) showed no differences among treatment groups. Additionally, the ratio of OCP/PO4 did not exhibit significant changes among treatment groups (D, H).

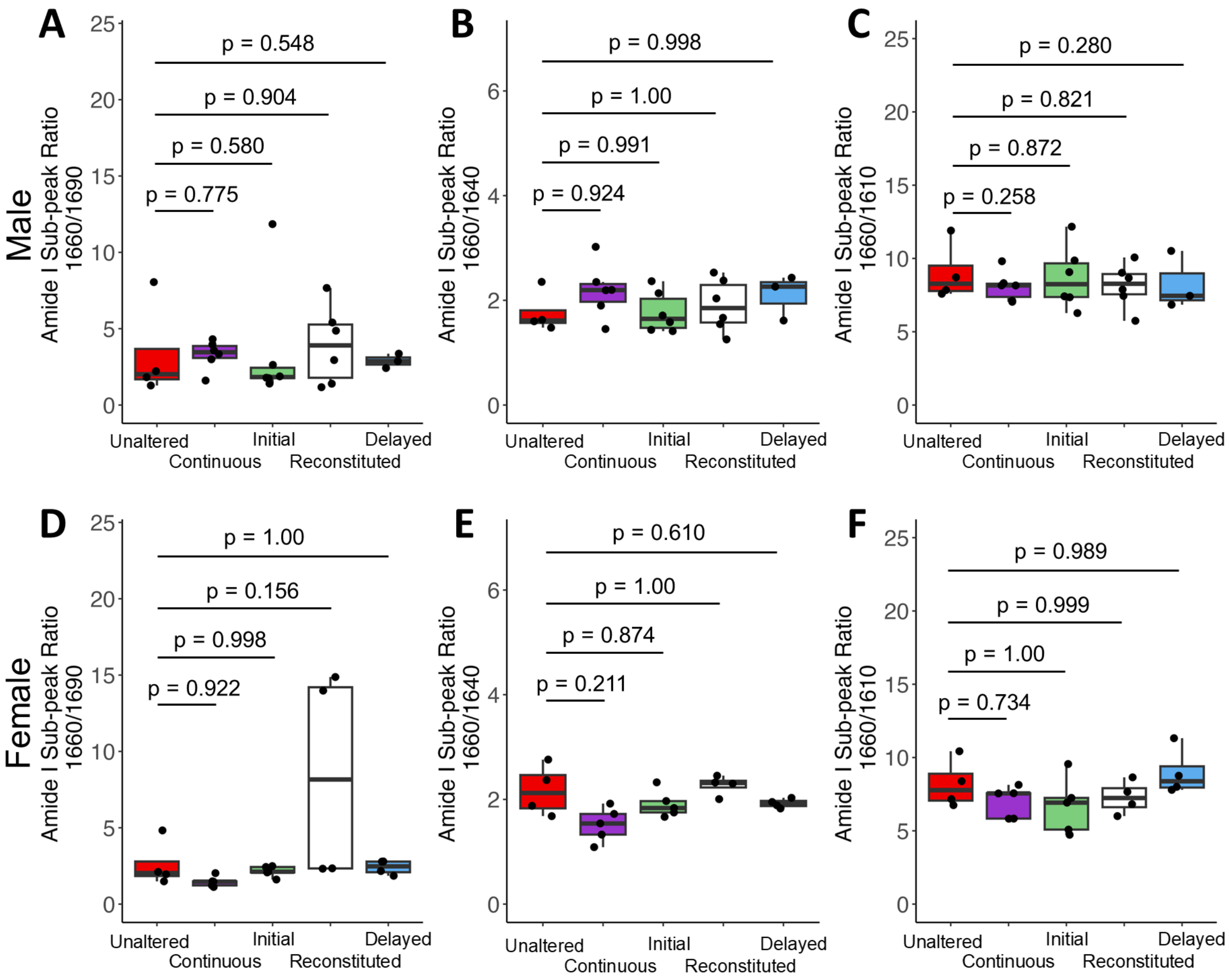


**Supplemental Figure 5.** Raman spectroscopy measurements of the amide I sub peak ratios. The 1660/1690 ratio measures the collagen maturity and there is no observed significance across treatment groups in both male and female (A, D). The 1660/1640 ratio measures the helical status of collagen and there is no observed significance across the treatment groups in both male and female (B, E). The ratio of 1660/1610 did not show significant changes across treatment groups in both male and female (C, F).

**Supplemental Table 3.** The matrix composition measured by Raman spectroscopy is correlated with the several physiological measurements in males. The 95% confidence interval is shown in the top right corner and the Pearson's correlation coefficient is shown in the bottom left corner. Coefficients in bold indicate p value less than 0.05.

| Male Raman and Physiological Correlations | | | | | | | | | | | | | | | | | | |
|---|---|---|---|---|---|---|---|---|---|---|---|---|---|---|---|---|---|---|
| | v1/A1 | v1/Pro | TBC | OCP/PO4 | v2/A3 | HPRO/PRO | CML | PEN | 1660/1640 | 1660/1690 | 1660/1610 | Crystallinity | Body Weight (g) | Max Moment (Nmm) | Cross-sectional Area (mm$^2$) | Section Modulus (mm$^3$) | Tissue Strength (MPa) | Work to Failure (N·mm) |
| v1/A1 | | [-0.047 0.529] | [0.217 0.694] | [0.024 0.584] | [0.228 0.700] | [-0.440 0.163] | [0.342 0.758] | [-0.203 0.521] | [-0.328 0.303] | [-0.314 0.309] | [-0.355 0.259] | [-0.362 0.251] | [-0.431 0.173] | [-0.116 0.491] | [-0.196 0.411] | [-0.187 0.419] | [-0.239 0.388] | [-0.565 0.013] |
| v1/Pro | 0.265 | | [-0.076 0.508] | [-0.234 0.385] | [-0.174 0.431] | [0.620 0.877] | [-0.202 0.406] | [-0.204 0.521] | [-0.298 0.333] | [-0.405 0.212] | [-0.521 0.058] | [-0.339 0.276] | [-0.247 0.366] | [-0.226 0.400] | [-0.191 0.415] | [-0.152 0.448] | [-0.305 0.326] | [-0.446 0.172] |
| TBC | **0.492** | 0.237 | | [-0.243 0.377] | [-0.142 0.457] | [-0.233 0.379] | [-0.125 0.470] | [-0.096 0.597] | [-0.240 0.388] | [-0.209 0.407] | [-0.333 0.282] | [-0.679 -0.189] | [-0.116 0.477] | [-0.312 0.319] | [0.046 0.593] | [0.161 0.663] | [-0.608 -0.052] | [-0.630 -0.088] |
| OCP/PO4 | **0.333** | 0.083 | 0.075 | | [-0.103 0.493] | [-0.417 0.198] | [-0.140 0.465] | [-0.475 0.261] | [-0.442 0.185] | [-0.451 0.166] | [-0.424 0.190] | [-0.277 0.345] | [-0.088 0.505] | [-0.004 0.577] | [0.033 0.590] | [-0.112 0.487] | [-0.197 0.432] | [-0.317 0.322] |
| v2/A3 | **0.501** | 0.142 | 0.173 | 0.215 | | [-0.349 0.265] | [-0.047 0.529] | [-0.520 0.204] | [-0.255 0.373] | [-0.305 0.318] | [-0.558 0.006] | [-0.426 0.179] | [-0.572 -0.014] | [-0.496 0.109] | [-0.370 0.243] | [-0.421 0.185] | [-0.435 0.185] | [-0.418 0.206] |
| HPRO/PRO | -0.153 | **0.779** | 0.081 | -0.121 | -0.046 | | [-0.478 0.115] | [-0.492 0.241] | [-0.278 0.352] | [-0.343 0.279] | [-0.365 0.248] | [-0.297 0.318] | [-0.133 0.464] | [-0.355 0.275] | [-0.156 0.445] | [-0.026 0.544] | [-0.510 0.090] | [-0.419 0.204] |
| CML | **0.588** | 0.113 | 0.190 | 0.179 | 0.264 | -0.199 | | [-0.063 0.618] | [-0.269 0.360] | [-0.148 0.458] | [-0.325 0.290] | [-0.299 0.316] | [-0.281 0.334] | [-0.051 0.539] | [-0.192 0.415] | [-0.234 0.378] | [-0.112 0.494] | [-0.274 0.356] |
| PEN | 0.184 | 0.183 | 0.288 | -0.1248 | -0.183 | -0.145 | 0.318 | | [-0.317 0.439] | [-0.313 0.444] | [-0.531 0.190] | [-0.402 0.344] | [-0.406 0.339] | [-0.295 0.472] | [-0.318 0.426] | [-0.349 0.396] | [-0.304 0.465] | [-0.360 0.414] |
| 1660/1640 | -0.014 | 0.020 | 0.082 | -0.143 | 0.066 | 0.041 | 0.050 | 0.071 | | [0.430 0.809] | [-0.279 0.351] | [-0.235 0.392] | [-0.624 - 0.079] | [-0.432 0.206] | [-0.434 0.187] | [-0.497 0.108] | [-0.290 0.357] | [-0.498 0.125] |
| 1660/1690 | -0.002 | -0.107 | 0.110 | -0.157 | 0.007 | -0.035 | 0.171 | 0.077 | **0.659** | | [0.066 0.611] | [-0.211 0.405] | [-0.474 0.128] | [-0.236 0.400] | [-0.242 0.378] | [-0.316 0.307] | [-0.237 0.397] | [-0.392 0.243] |
| 1660/1610 | -0.053 | -0.254 | -0.028 | -0.129 | -0.302 | -0.065 | -0.019 | -0.197 | 0.040 | **0.3702** | | [-0.099 0.490] | [-0.246 0.367] | [-0.246 0.382] | [-0.219 0.392] | [-0.111 0.481] | [-0.415 0.209] | [-0.340 0.290] |
| Crystallinity | -0.061 | -0.035 | **-0.469** | 0.038 | -0.136 | 0.011 | 0.009 | -0.034 | 0.087 | 0.107 | 0.215 | | [-0.403 0.206] | [-0.212 0.412] | [-0.405 0.204] | [-0.422 0.184] | [-0.111 0.495] | [-0.075 0.522] |
| Body Weight (g) | -0.142 | 0.066 | 0.200 | 0.230 | **-0.320** | 0.182 | 0.02 | -0.039 | **-0.385** | -0.191 | 0.067 | -0.109 | | [0.150 0.553] | [0.337 0.677] | [0.526 0.785] | [-0.351 0.105] | [-0.151 0.309] |
| Max Moment (Nmm) | 0.207 | 0.097 | 0.003 | 0.316 | -0.214 | -0.044 | 0.269 | 0.104 | -0.126 | 0.091 | 0.076 | 0.111 | **0.369** | | [0.202 0.589] | [0.147 0.551] | [0.617 0.831] | [0.394 0.710] |
| Cross-sectional Area (mm$^2$) | 0.119 | 0.124 | **0.348** | **0.341** | -0.070 | 0.160 | 0.123 | 0.062 | -0.137 | 0.075 | 0.095 | -0.111 | **0.528** | **0.414** | | [0.830 0.930] | [-0.416 0.029] | [-0.196 0.267] |
| Section Modulus (mm$^3$) | 0.128 | 0.163 | **0.447** | 0.207 | -0.130 | 0.284 | 0.079 | 0.027 | -0.215 | -0.005 | 0.204 | -0.132 | **0.676** | **0.366** | **0.891** | | [-0.527 -0.113] | [-0.197 0.266] |
| Tissue Strength (MPa) | 0.082 | 0.012 | **-0.362** | 0.131 | -0.139 | -0.232 | 0.211 | 0.095 | 0.037 | 0.089 | -0.114 | 0.212 | -0.130 | **0.742** | -0.204 | **-0.336** | | [0.349 0.684] |
| Work to Failure (N·mm) | -0.304 | -0.153 | **-0.393** | 0.003 | -0.118 | -0.119 | 0.046 | 0.031 | -0.208 | -0.083 | -0.028 | 0.247 | 0.083 | **0.573** | 0.038 | 0.036 | **0.537** | |

**Supplemental Table 4.** The matrix composition measured by Raman spectroscopy is correlated with the several physiological measurements in females. The 95% confidence interval is shown in the top right corner and the Pearson's correlation coefficient is shown in the bottom left corner. Coefficients in bold indicate p value less than 0.05.

| Female Raman and Physiological Correlations | | | | | | | | | | | | | | | | | | |
|---|---|---|---|---|---|---|---|---|---|---|---|---|---|---|---|---|---|---|
| | v1/A1 | v1/Pro | TBC | OCP/PO4 | v2/A3 | HPRO/PRO | CML | PEN | 1660/1640 | 1660/1690 | 1660/1610 | Crystallinity | Body Weight (g) | Max Moment (Nmm) | Cross-sectional Area (mm$^2$) | Section Modulus (mm$^3$) | Tissue Strength (MPa) | Work to Failure (N·mm) |
| v1/A1 | | [0.248 0.703] | [0.090 0.616] | [-0.293 0.307] | [0.183 0.671] | [-0.521 0.0579] | [0.213 0.688] | [-0.119 0.561] | [-0.419 0.179] | [-0.371 0.249] | [-0.401 0.200] | [-0.547 0.006] | [-0.402 0.192] | [-0.475 0.208] | [-0.500 0.095] | [-0.520 0.068] | [-0.430 0.262] | [-0.695 - 0.129] |
| v1/Pro | **0.510** | | [-0.040 0.517] | [-0.125 0.446] | [-0.150 0.437] | [-0.328 0.272] | [-0.069 0.495] | [-0.190 0.490] | [-0.306 0.287] | [-0.384 0.219] | [-0.359 0.232] | [-0.474 0.089] | [-0.334 0.252] | [-0.279 0.395] | [-0.489 0.093] | [-0.336 0.271] | [-0.337 0.339] | [-0.631 - 0.038] |
| TBC | **0.384** | 0.260 | | [-0.235 0.357] | [-0.075 0.502] | [-0.434 0.162] | [0.144 0.644] | [-0.076 0.581] | [-0.221 0.376] | [-0.554 0.011] | [-0.351 0.248] | [-0.793 -0.437] | [-0.403 0.183] | [-0.208 0.466] | [-0.445 0.156] | [-0.521 0.059] | [-0.135 0.523] | [-0.750 - 0.251] |
| OCP/PO4 | 0.008 | 0.175 | 0.067 | | [-0.341 0.259] | [-0.329 0.271] | [-0.078 0.489] | [0.185 0.723] | [-0.475 0.096] | [-0.366 0.240] | [-0.399 0.188] | [-0.394 0.186] | [-0.465 0.101] | [-0.372 0.304] | [-0.261 0.346] | [-0.103 0.481] | [-0.470 0.191] | [-0.354 0.322] |
| v2/A3 | **0.461** | 0.158 | 0.234 | -0.045 | | [-0.606 - 0.067] | [-0.201 0.400] | [0.004 0.632] | [-0.42 0.178] | [-0.387 0.232] | [-0.240 0.365] | [-0.604 -0.079] | [-0.300 0.300] | [-0.379 0.307] | [-0.355 0.267] | [-0.395 0.223] | [-0.358 0.328] | [-0.591 0.036] |
| HPRO/PRO | -0.254 | -0.031 | -0.150 | -0.032 | **-0.367** | | [-0.422 0.176] | [-0.583 0.087] | [-0.425 0.173] | [-0.489 0.110] | [-0.350 0.256] | [-0.122 0.460] | [-0.389 0.206] | [-0.296 0.389] | [-0.438 0.173] | [-0.394 0.224] | [-0.235 0.443] | [-0.225 0.451] |
| CML | **0.485** | 0.233 | 0.426 | 0.224 | 0.109 | -0.135 | | [-0.257 0.444] | [-0.491 0.083] | [-0.402 0.206] | [-0.308 0.293] | [-0.433 0.147] | [-0.353 0.239] | [-0.246 0.434] | [-0.411 0.197] | [-0.355 0.258] | [-0.221 0.455] | [-0.576 0.059] |
| PEN | 0.252 | 0.170 | 0.286 | **0.501** | **0.358** | -0.282 | 0.107 | | [-0.541 0.134] | [-0.470 0.227] | [-0.553 0.117] | [-0.501 0.175] | [-0.423 0.270] | [-0.306 0.463] | [-0.373 0.324] | [-0.347 0.350] | [-0.339 0.433] | [-0.530 0.223] |
| 1660/1640 | -0.132 | -0.010 | 0.08 | -0.207 | -0.133 | -0.139 | -0.223 | -0.232 | | [-0.033 0.539] | [-0.377 0.220] | [-0.351 0.241] | [-0.106 0.467] | [-0.177 0.491] | [-0.263 0.351] | [-0.098 0.491] | [-0.317 0.369] | [-0.238 0.441] |
| 1660/1690 | -0.067 | -0.091 | *-0.297* | -0.069 | -0.086 | -0.209 | -0.108 | -0.139 | 0.277 | | [-0.338 0.277] | [-0.037 0.530] | [-0.419 0.180] | [-0.320 0.377] | [-0.334 0.297] | [-0.153 0.462] | [-0.407 0.288] | [0.156 0.709] |
| 1660/1610 | -0.111 | -0.070 | -0.056 | -0.115 | 0.069 | -0.052 | -0.008 | -0.248 | -0.086 | -0.034 | | [-0.246 0.347] | [-0.243 0.349] | [-0.675 - 0.104] | [0.218 0.695] | [0.022 0.577] | [-0.729 - 0.208] | [-0.607 0.011] |
| Crystallinity | *-0.295* | -0.210 | **-0.649** | -0.114 | **-0.371** | 0.185 | -0.156 | -0.185 | -0.060 | 0.270 | 0.055 | | [-0.403 0.206] | [-0.211 0.412] | [-0.405 0.204] | [-0.422 0.184] | [-0.111 0.495] | [-0.075 0.522] |
| Weight (g) | -0.115 | -0.045 | -0.120 | -0.198 | 0.000 | -0.101 | -0.062 | -0.087 | 0.197 | -0.132 | 0.058 | 0.230 | | [0.131 0.572] | [0.353 0.710] | [0.330 0.700] | [-0.226 0.281] | [-0.128 0.371] |
| Max Moment (Nmm) | -0.151 | 0.066 | 0.146 | -0.038 | -0.041 | 0.053 | 0.107 | 0.092 | 0.177 | 0.032 | **-0.432** | 0.117 | **0.373** | | [0.091 0.544] | [-0.035 0.450] | [0.710 0.887] | [0.143 0.581] |
| Cross-sectional Area (mm$^2$) | -0.223 | -0.217 | -0.160 | 0.047 | -0.048 | -0.147 | -0.118 | -0.028 | 0.049 | -0.020 | **0.492** | 0.195 | **0.557** | **0.337** | | [0.919 0.971] | [-0.508 -0.040] | [-0.178 0.327] |
| Section Modulus (mm$^3$) | -0.249 | -0.036 | -0.254 | 0.207 | -0.095 | -0.094 | -0.053 | 0.001 | 0.216 | 0.171 | **0.327** | **0.313** | **0.538** | 0.221 | **0.869** | | [-0.575 -0.136] | [-0.147 0.355] |
| Tissue Strength (MPa) | -0.095 | 0.001 | 0.219 | -0.157 | -0.017 | 0.118 | 0.132 | 0.055 | 0.029 | -0.068 | **-0.514** | 0.010 | 0.030 | **0.816** | -0.177 | **-0.376** | | [0.033 0.502] |
| Work to Failure (N·mm) | **-0.457** | **-0.372** | **-0.547** | -0.019 | -0.311 | 0.128 | -0.290 | -0.180 | 0.115 | **0.479** | *-0.333* | **0.479** | 0.130 | **0.383** | 0.025 | 0.111 | **0.285** | |

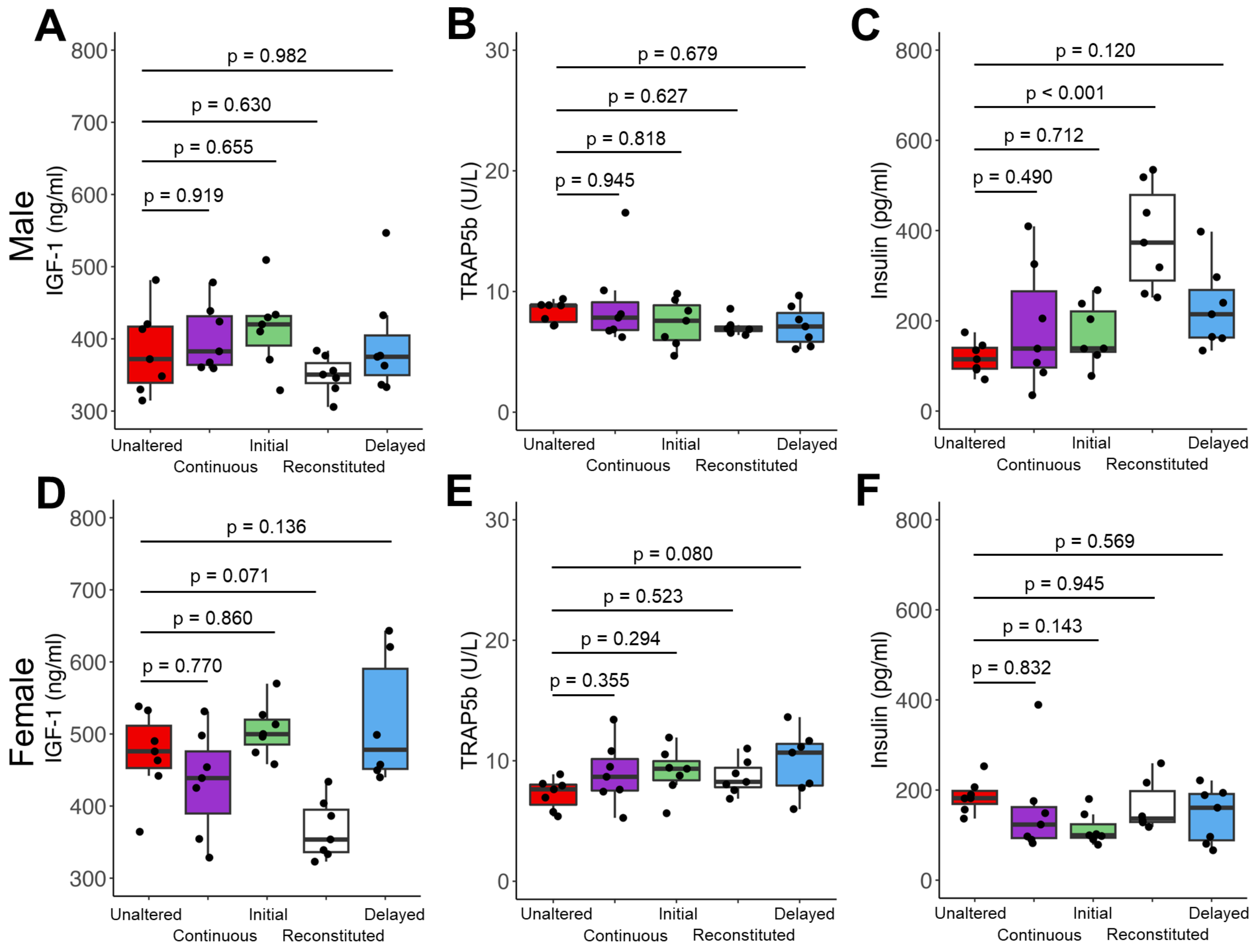


**Supplemental Figure 6.** The serum concentration of the bone resorption marker TRAP5b did not show significant differences among treatment groups (B, E). Similarly, there were no significant differences in the growth hormone IGF-1 among treatment groups (A, D). The Reconstituted male showed an increase in the insulin level compared to the Untreated (C) but not in females (F).

**Supplemental Table 5**. The serum markers are correlated with the several physiological measurements in males. The 95% confidence interval is shown in the top right corner and the Pearson’s correlation coefficient is shown in the bottom left corner. Coefficients in bold indicate p value less than 0.05.

| Male Serum and Physiological Correlations | | | | | | | | | | | | | | | |
|---|---|---|---|---|---|---|---|---|---|---|---|---|---|---|---|
| | IL-6 (pg/ml) | Insulin (pg/ml) | Leptin (pg/ml) | TNF-α (pg/ml) | Adiponectin (pg/ml) | IGF-1 (pg/ml) | P1NP (ng/ml) | TRAP5b (U/L) | Femur Length (mm) | Maximum Moment (N·mm) | Body Weight (g) | Cross-sectional Area (mm$^2$) | Section Modulus (mm$^3$) | Tissue Strength (MPa) | Work to Failure (N·mm) |
| IL-6 (pg/ml) | | [-0.554 0.115] | [-0.485 0.208] | [0.240 0.756] | [-0.585 0.071] | [-0.474 0.222] | [-0.414 0.292] | [-0.201 0.501] | [0.037 0.651] | [-0.439 0.263] | [-0.196 0.495] | [-0.365 0.343] | [-0.273 0.430] | [-0.520 0.163] | [-0.263 0.440] |
| Insulin (pg/ml) | -0.249 | | [-0.241 0.420] | [-0.800 -0.380] | [-0.508 0.132] | [-0.629 -0.046] | [-0.715 -0.201] | [-0.507 0.144] | [-0.259 0.404] | [-0.243 0.417] | [0.027 0.617] | [-0.361 0.305] | [-0.304 0.362] | [-0.225 0.433] | [-0.244 0.417] |
| Leptin (pg/ml) | -0.158 | 0.101 | | [-0.365 0.300] | [-0.192 0.461] | [-0.171 0.478] | [-0.206 0.449] | [-0.225 0.442] | [-0.248 0.414] | [-0.293 0.373] | [-0.326 0.340] | [-0.440 0.217] | [-0.424 0.236] | [-0.224 0.434] | [-0.357 0.309] |
| TNF-α (pg/ml) | **0.548** | **-0.633** | -0.037 | | [0.169 0.479] | [0.105 0.663] | [0.145 0.685] | [0.012 0.614] | [-0.153 0.492] | [-0.628 -0.046] | [-0.598 0.003] | [-0.430 0.229] | [-0.434 0.224] | [-0.648 -0.078] | [-0.573 0.040] |
| Adiponectin (pg/ml) | -0.291 | -0.210 | 0.151 | 0.174 | | [-0.036 0.576] | [-0.027 0.582] | [-0.467 0.195] | [-0.436 0.222] | [-0.447 0.209] | [-0.652 -0.086] | [-0.538 0.091] | [-0.544 0.083] | [-0.301 0.364] | [-0.381 0.284] |
| IGF-1 (pg/ml) | -0.144 | **-0.374** | 0.172 | **0.424** | 0.301 | | [0.160 0.693] | [-0.097 0.542] | [-0.445 0.211] | [-0.468 0.183] | [-0.522 0.113] | [-0.410 0.252] | [-0.420 0.240] | [-0.444 0.213] | [-0.463 0.189] |
| P1NP (ng/ml) | -0.069 | **-0.501** | 0.136 | **0.456** | 0.309 | **0.468** | | [-0.087 0.549] | [-0.326 0.340] | [-0.404 0.258] | [-0.513 0.125] | [-0.317 0.349] | [-0.321 0.346] | [-0.451 0.204] | [-0.413 0.249] |
| TRAP5b (U/L) | 0.171 | -0.204 | 0.122 | **0.349** | -0.153 | 0.249 | 0.259 | | [-0.092 0.545] | [-0.207 0.458] | [-0.180 0.480] | [-0.111 0.532] | [-0.036 0.584] | [-0.390 0.285] | [-0.238 0.431] |
| Femur Length (mm) | **0.386** | 0.081 | 0.093 | 0.190 | -0.120 | -0.131 | 0.008 | 0.254 | | [-0.134 0.325] | [0.302 0.655] | [0.162 0.561] | [0.209 0.594] | [-0.425 0.018] | [-0.371 0.082] |
| Maximum Moment (N·mm) | -0.100 | 0.098 | 0.045 | **-0.373** | -0.134 | -0.160 | -0.082 | 0.142 | 0.101 | | [0.150 0.553] | [0.202 0.589] | [0.147 0.551] | [0.617 0.831] | [0.394 0.710] |
| Body Weight (g) | 0.171 | **0.357** | 0.008 | -0.330 | **-0.407** | -0.229 | -0.217 | 0.169 | **0.499** | **0.369** | | [0.337 0.677] | [0.526 0.785] | [-0.351 0.105] | [-0.151 0.309] |
| Cross-sectional Area (mm$^2$) | -0.012 | -0.032 | -0.126 | -0.113 | -0.250 | -0.089 | 0.018 | 0.236 | **0.379** | **0.414** | **0.528** | | [0.830 0.930] | [-0.416 0.029] | [-0.196 0.267] |
| Section Modulus (mm$^3$) | 0.090 | 0.032 | -0.106 | -0.118 | -0.257 | -0.101 | 0.014 | 0.306 | **0.420** | **0.366** | **0.676** | **0.891** | | [-0.527 -0.113] | [-0.197 0.266] |
| Tissue Strength (MPa) | -0.203 | 0.117 | 0.118 | **-0.401** | 0.035 | -0.130 | -0.138 | -0.059 | -0.214 | **0.742** | -0.130 | -0.204 | **-0.336** | | [0.349 0.684] |
| Work to Failure (N·mm) | 0.101 | 0.097 | -0.027 | -0.297 | -0.055 | -0.154 | -0.092 | 0.108 | -0.152 | **0.573** | 0.083 | 0.038 | 0.036 | **0.537** | |

**Supplemental Table 6.** The serum markers are correlated with the several physiological measurements in females. The 95% confidence interval is shown in the top right corner and the Pearson's correlation coefficient is shown in the bottom left corner. Coefficients in bold indicate p value less than 0.05.

| Female Serum and Physiological Correlations | | | | | | | | | | | | | | | |
|---|---|---|---|---|---|---|---|---|---|---|---|---|---|---|---|
| | IL-6 (pg/ml) | Insulin (pg/ml) | Leptin (pg/ml) | TNF-α (pg/ml) | Adiponectin (pg/ml) | IGF-1 (pg/ml) | P1NP (ng/ml) | TRAP5b (U/L) | Femur Length (mm) | Maximum Moment (N·mm) | Body Weight (g) | Cross-sectional Area ($mm^2$) | Section Modulus ($mm^3$) | Tissue Strength (MPa) | Work to Failure (N·mm) |
| IL-6 (pg/ml) | | [-0.449 0.278] | [-0.588 0.093] | [-0.093 0.598] | [-0.363 0.370] | [-0.444 0.283] | [-0.269 0.456] | [-0.189 0.521] | [-0.396 0.337] | [-0.623 0.038] | [-0.409 0.323] | [-0.431 0.298] | [-0.451 0.275] | [-0.574 0.115] | [-0.697 -0.092] |
| Insulin (pg/ml) | -0.099 | | [-0.200 0.463] | [-0.357 0.330] | [-0.465 0.197] | [-0.258 0.414] | [-0.523 0.123] | [-0.548 0.088] | [-0.572 0.077] | [-0.231 0.456] | [-0.101 0.556] | [-0.243 0.446] | [-0.159 0.514] | [-0.352 0.345] | [-0.286 0.409] |
| Leptin (pg/ml) | -0.283 | 0.148 | | [-0.567 0.073] | [0.209 0.724] | [-0.403 0.270] | [-0.606 0.001] | [-0.491 0.164] | [-0.269 0.423] | [0.132 0.696] | [0.346 0.796] | [-0.189 0.491] | [-0.206 0.477] | [-0.001 0.621] | [-0.138 0.529] |
| TNF-α (pg/ml) | 0.290 | -0.015 | -0.277 | | [-0.088 0.548] | [0.144 0.690] | [0.156 0.697] | [-0.237 0.432] | [-0.608 0.035] | [-0.444 0.258] | [-0.613 0.027] | [-0.281 0.424] | [-0.304 0.402] | [-0.458 0.241] | [-0.691 -0.108] |
| Adiponectin (pg/ml) | 0.004 | -0.151 | 0.511 | 0.258 | | [-0.359 0.307] | [-0.160 0.490] | [-0.359 0.307] | [-0.416 0.278] | [-0.081 0.570] | [0.050 0.652] | [-0.200 0.482] | [-0.279 0.415] | [-0.138 0.529] | [-0.480 0.202] |
| IGF-1 (pg/ml) | -0.093 | 0.088 | -0.075 | **0.460** | 0.083 | | [-0.374 0.292] | [-0.290 0.375] | [-0.205 0.477] | [-0.301 0.395] | [-0.505 0.170] | [-0.206 0.477] | [-0.314 0.382] | [-0.326 0.371] | [-0.513 0.160] |
| P1NP (ng/ml) | 0.108 | -0.224 | -0.337 | **0.470** | 0.183 | -0.046 | | [-0.249 0.412] | [-0.439 0.251] | [-0.447 0.242] | [-0.629 -0.011] | [-0.486 0.195] | [-0.439 0.252] | [-0.395 0.300] | [-0.476 0.208] |
| TRAP5b (U/L) | 0.191 | -0.258 | -0.184 | 0.110 | -0.029 | 0.048 | 0.091 | | [-0.608 0.023] | [-0.534 0.131] | [-0.695 -0.129] | [-0.502 0.174] | [-0.611 0.017] | [-0.374 0.323] | [-0.539 0.125] |
| Femur Length (mm) | -0.034 | -0.279 | 0.087 | -0.324 | -0.078 | 0.154 | -0.107 | -0.329 | | [0.073 0.532] | [0.301 0.681] | [0.235 0.640] | [0.226 0.635] | [-0.221 0.286] | [0.056 0.519] |
| Maximum Moment (N·mm) | -0.333 | 0.128 | **0.459** | -0.106 | 0.276 | 0.053 | -0.117 | -0.228 | **0.321** | | [0.131 0.572] | [0.091 0.544] | [-0.035 0.450] | [0.710 0.887] | [0.143 0.581] |
| Body Weight (g) | -0.050 | 0.257 | **0.620** | -0.330 | **0.392** | -0.189 | **-0.358** | **-0.457** | **0.516** | **0.373** | | [0.091 0.544] | [-0.035 0.450] | [0.710 0.887] | [0.143 0.581] |
| Cross-sectional Area ($mm^2$) | -0.077 | 0.116 | 0.171 | 0.082 | 0.160 | 0.153 | -0.165 | -0.186 | **0.461** | **0.337** | **0.557** | | [0.919 0.971] | [-0.508 -0.040] | [-0.178 0.327] |
| Section Modulus ($mm^3$) | -0.101 | 0.201 | 0.154 | 0.056 | 0.077 | 0.038 | -0.106 | -0.334 | **0.454** | 0.221 | **0.538** | **0.869** | | [-0.575 -0.136] | [-0.147 0.355] |
| Tissue Strength (MPa) | -0.262 | -0.004 | 0.348 | -0.124 | 0.221 | 0.026 | -0.054 | -0.029 | 0.035 | **0.816** | 0.030 | -0.177 | **-0.376** | | [0.033 0.502] |
| Work to Failure (N·mm) | **-0.443** | 0.070 | 0.221 | **-0.445** | -0.158 | -0.200 | -0.152 | -0.234 | **0.305** | **0.383** | 0.130 | 0.025 | 0.111 | **0.285** | |

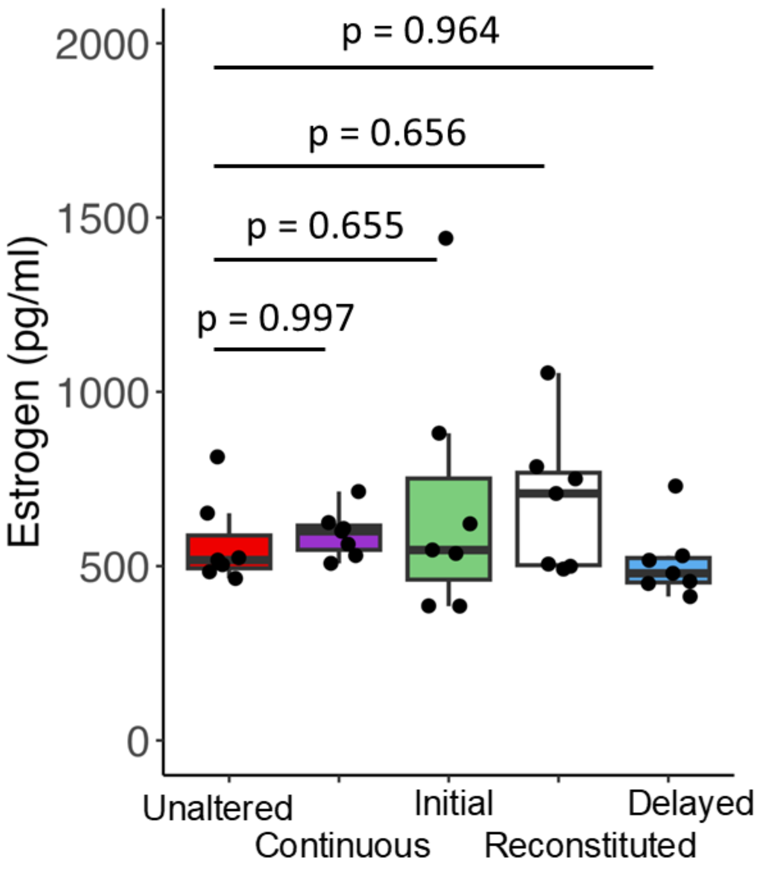


**Supplemental Figure 6.** The serum concentration of estrogen showed no differences among the treatment groups in females.
